# NudC regulated Lis1 stability is essential for maintenance of dynamic microtubule ends in the axon terminal

**DOI:** 10.1101/2022.07.12.499810

**Authors:** Dane Kawano, Katherine Pinter, Madison Chlebowski, Ronald S. Petralia, Ya-Xian Wang, Alex V. Nechiporuk, Catherine M Drerup

**Affiliations:** Unit on Neuronal Cell Biology, National Institute of Child Health & Human Development, National Institutes of Health, Bethesda, MD 20892; Advanced Imaging Core, National Institute of Deafness and other Communication Disorders, National Institutes of Health, Bethesda, MD 20892; Department of Cell, Developmental & Cancer Biology, Oregon Health & Science University, Portland, OR 97239; Department of Integrative Biology, University of Wisconsin-Madison, Madison, WI 53706

## Abstract

Axon terminal structure is critical for neuronal function. This cellular compartment houses synaptic terminals and is a site of high metabolic and functional demand. Axon terminals are also the site of a change in microtubule structure within the neuron. Microtubule stability is decreased relative to the axon shaft due to an enrichment of microtubule plus ends and increase in microtubule dynamics. These dynamic microtubule plus ends have many functions including serving as a docking site for the microtubule motor protein complex Cytoplasmic dynein. Here, we report an unexplored function of the dynein motor in axon terminals: regulation of microtubule stability. Using a forward genetic screen, we identified a mutant with abnormal axon terminal structure due to a loss of function mutation in the dynein interacting protein NudC. We show that the primary function of NudC in the axon terminal is as a chaperone for the protein Lis1. Loss of NudC results in decreased Lis1 protein in this neuronal compartment. Decreased Lis1 in *nudc* mutants causes dynein/dynactin accumulation and increased microtubule stability in axon terminals. Microtubules in the proximal axon are unaffected. Abnormal microtubule stability and structure can be suppressed by pharmacologically inhibiting dynein, implicating excess dynein motor activity as causal in the enhanced axon terminal microtubule stability. Together, our data support a model in which local NudC-Lis1 modulation of dynein motor activity is critical for regulation of microtubule stability in the axon terminal.

## Introduction

In neurons, microtubules have varying levels of stability based on the cellular compartment. In the axon, microtubules along the axon shaft are relatively stable compared to those in the axon terminal [1]. The maintenance of these dynamic microtubule plus ends is critical to neuron structure and function. One role for the microtubule plus ends in the axon terminal is docking of the motor protein complex Cytoplasmic dynein (referred to as dynein hereafter). Dynein docks on microtubule plus ends prior to cargo binding and initiation of retrograde transport. Docking of dynein on these plus ends has been shown to rely on dynactin interaction with microtubule plus end binding proteins [2–4]. Disrupting this process inhibits initiation of retrograde cargo transport [2, 5, 6].

Dynein works with adaptor proteins to bind vesicular cargo to the motor for retrograde transport from microtubule plus ends [7–9]. In addition to adaptor proteins, Lis1 (Lissencephaly 1) has received significant attention for its role in facilitating dynein-cargo interaction. Structural and in vivo evidence suggests that Lis1 can facilitate dynein interaction with activating adaptors and, subsequently, incorporation of paired dynein motor dimers with cargo for efficient retrograde transport [10–12]. This is likely through changing the propensity for dynein to be in an “open” conformation which encourages adaptor interaction [12, 13]. The combined efforts of Lis1 and activating adaptors are essential for efficient retrograde transport from axon terminals.

Lis1 and dynein have also been implicated in regulation of microtubule stability, though whether they increase or decrease microtubule stability differs based on cell type and model. In vitro, in the presence of ATP, dynein has been shown to bind to the microtubule lattice and stabilize it. This prevents collapse typically induced when a growing microtubule encounters a barrier [14]. In vivo, loss of dynein function leads to microtubule instability in the tips of dendritic processes in *C. elegans* [15]. Conversely, in Aspergillus and budding yeast, there is evidence that dynein destabilizes microtubules, particularly during microtubule contact with the cell cortex [16–19]. Similarly, whether Lis1 increases or decreases stability is controversial and may depend upon context or cell type [17, 18, 20–22]. While we have learned a great deal about the activators required for dynein-dependent retrograde transport initiation, an in-depth understanding of dynein-dependent microtubule regulation in axons is still lacking.

We used a forward genetic screen in zebrafish to identify novel dynein modulators in neurons. For this screen and subsequent work, we took advantage of the sensory axons of the zebrafish posterior lateral line (pLL) system. The long axons of the pLL extend from the pLL ganglion (pLLg) just behind the ear to the tip of the tail. These axons fully extend by 2 days post fertilization (dpf) and by 4 dpf the axons of the pLL have made active synaptic contacts with their target sensory cells. The pLL axons and their associated sensory organs are superficial, allowing for direct in vivo observation of axon morphology and axonal transport dynamics in a fully intact animal (Figure 1A).

**Figure 1.**
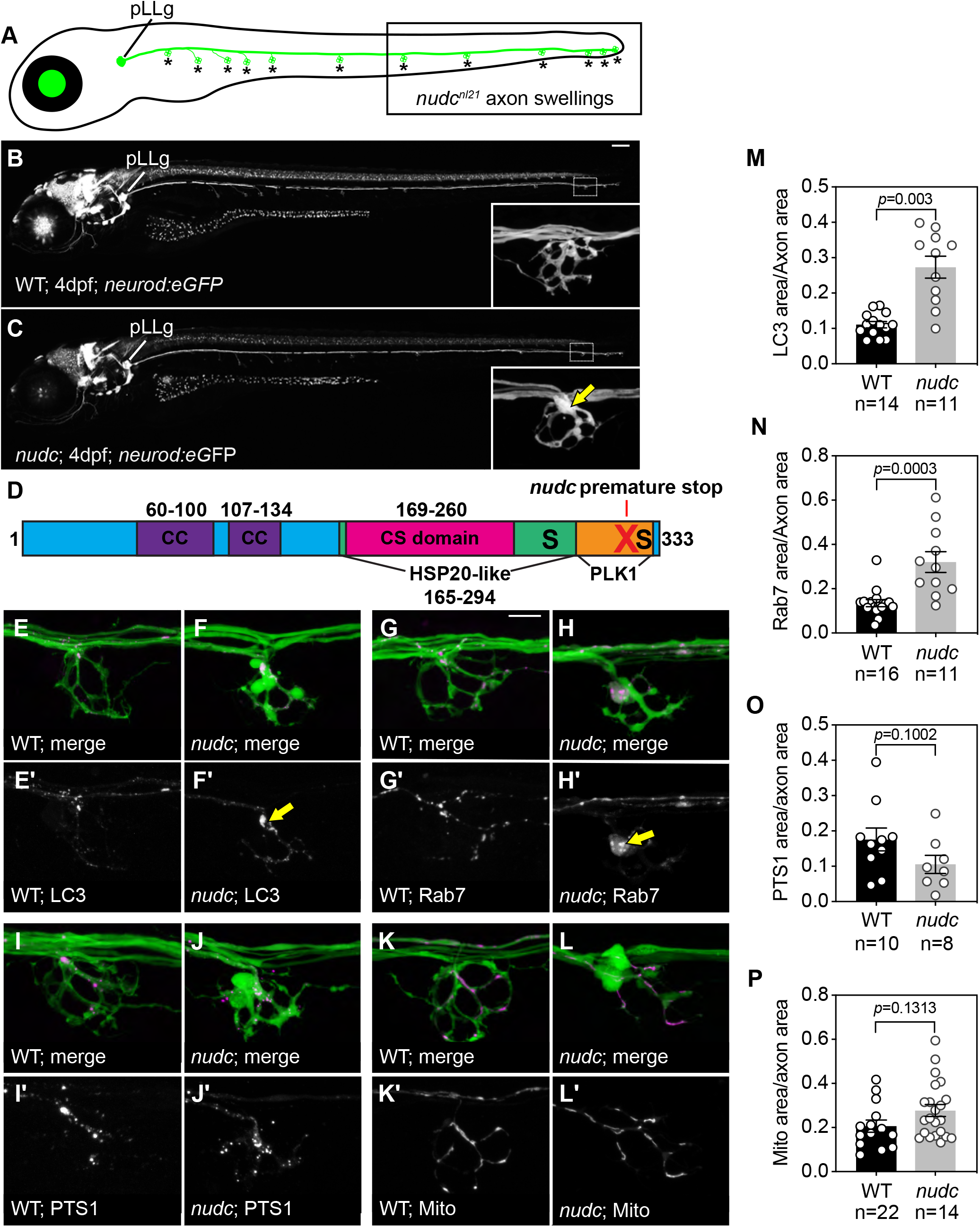
NudC loss of function causes axon terminal swellings in the long axons of the zebrafish posterior lateral line (pLL). (A) Schematic of a larval zebrafish at 4 dpf. The pLL ganglion (pLLg) is indicated. Asterisks point to axon terminals in the pLL system. The posterior half of the trunk is where pLL axon terminal swellings are clearly observed in *nudc* mutants (boxed). (B,C) Wild type and *nudc TgBAC*(*neurod:egfp*)*^nl1^* larva at 4 dpf. Insets show magnified view of a distal axon terminal. Arrow points to axon terminal swelling in *nudc* mutants. (D) Diagram of NudC domains, including Plk1 phosphorylation sites at S274 and S326 (S) and the premature stop site in the *nudc^nl21^* line (red X). Also shown are the coiled coil (CC), HSP20-like, and CS domains. (E-L) Localization of autophagosomes (E,F), late endosomes (G,H), peroxisomes (I,J), and mitochondria (K,L) in wild type and *nudc* axon terminals. Arrows in (F’) and (H’) indicate accumulation of LC3+ and Rab7+ vesicles in nudc mutant axon terminals. (M-P) Average organelle load (organelle area/axon area) in axon terminals (Wilcoxon rank sum). Data are expressed as mean±S.E.M. Sample sizes indicated. Scale bar=100μm in B,C; 10μm in E-L.

From this screen, we identified a zebrafish strain with a loss of function mutation in *nudc* (*nudc^nl21^*). NudC was originally identified as a modulator of cell wall thickness and nuclear positioning through its association with Lis1 (also known as NudF) in the filamentous fungus *Aspergillus nidulans* [23–25]. NudC has also been implicated in dynein-dynactin function during cell migration in the developing cortex [26–28]. Finally, NudC has a conserved CS domain predicted to give it chaperone activity [29]. This chaperone function stabilizes proteins, including the dynein regulator Lis1 [24, 30]. Beyond this, its function in post-mitotic neurons and, more specifically, its regulation of microtubule motor-dependent cellular processes in axons are not well understood.

Using live imaging of organelle localization and transport as well as microtubule dynamics in zebrafish pLL axons, we show that NudC regulates dynein function by increasing Lis1 protein stability in the axon terminal. In *nudc* mutants, axon terminals are swollen with accumulations of dynein/dynactin components and common retrograde cargos. In addition, *nudc* mutant axon terminals contain nets of stabilized microtubules. These defects are due to loss of Lis1 in *nudc* mutants as exogenous expression of Lis1 can rescue both cargo accumulation and microtubule stability phenotypes. Increased levels of active dynein in the axon terminal appear to directly cause increased microtubule stabilization as pharmacological suppression of dynein activity can rescue microtubule structure. Together, our data suggest a model in which loss of NudC, and subsequently Lis1, leads to failed dynein-cargo attachment for initiation of retrograde transport. This results in an accumulation of not only cargo but also dynein in the axon terminal which alters microtubule stability in this neuronal compartment.

## Results

### A C-terminal mutation in *nudc* causes autophagosome accumulation in axon terminals

We undertook an ENU-induced, forward genetic screen in zebrafish to identify modulators of dynein in axons. This screen has been described in detail previously [31] and was used to isolated the *nudc^nl21^* mutant strain (hereafter referred to as *nudc*). The *nudc* mutant was identified due to the presence of axonal swellings at the branchpoints of the long axons of the pLL nerve (Figure 1A-C). The *nudc* mutation is a recessive T to A point mutation at the splice donor site of intron 8. This mutation leads to inclusion of intron 8 and a premature stop codon immediately following the exon (Figure 1D). Though close to the C-terminus of the protein, the mutation results in a loss of NudC function as the mutant axon terminal swelling phenotype can be suppressed by exogenous expression of wildtype NudC (Figure S1A).

Homozygous *nudc* mutants develop normally during larval stages but are not viable to adulthood. This is despite the fact that NudC is essential for cell division [32]. We reasoned this was due to the maternal deposition of wildtype *nudc* mRNA from heterozygous females as all mutants are derived from heterozygous crosses. To test this, we performed RT-PCR analysis from 2-cell stage embryos which confirmed maternal deposition of *nudc* mRNA prior to the onset of zygotic transcription (Figure S1B). This likely leads to sufficient NudC protein levels for cell division into larval stages; a similar maternal deposition of p150 mRNA leads to survival of *p150a* mutants in which this dynactin protein is lost [33]. Previous mutants identified from our screen with similar axonal phenotypes were shown to have disruptions to proteins critical for dynein function in axons [31, 34]. NudC interacts with dynein during cell division and during interkinetic nuclear migration in the developing cortex [26, 27]. In *Aspergillus* and cultured cells, NudC regulates the stability of the dynein activator Lis1 [24, 30, 35]. A role for NudC in the axon terminal had not been explored. First, we asked if the axon terminal swelling phenotype in *nudc* mutants was due to accumulations of any common dynein cargos. We addressed this using transmission electron microscopy (TEM) and in vivo imaging. TEM analyses revealed enlarged, double membrane bound compartments in *nudc* mutant axon terminals reminiscent of enlarged autophagosomes (Figure S1C,D). Quantification of autophagosome number showed an increase in *nudc* mutant TEMs (Figure S1E). To quantify total autophagosome density in axon terminals, we used live imaging to analyze the localization of autophagosomes by expressing RFP-tagged LC3 in pLL neurons. Images and quantification of LC3+ autophagosome density in axon terminals demonstrated a strong increase in *nudc* mutants (Figure 1E,F,M).

Late endosomes are typical cargo engulfed during bulk autophagy. In our TEM analyses, we observed small vesicles inside the enlarged autophagosomes and reasoned these could be late endosomes. To determine if late endosomes also accumulate in *nudc* mutant axon terminals, we analyzed the localization of a late endosome marker, Rab7. Live imaging revealed accumulations of Rab7+ late endosomes in *nudc* mutant axon terminals (Figure 1G,H,N). Finally, we asked if all retrograde cargos were accumulating in *nudc* mutant axon terminals. To address this, we assayed the localization of cargos largely unrelated to autophagosomes, peroxisomes and mitochondria. Live imaging of axon terminals revealed no change in peroxisome or mitochondrial density in the axon terminal (Figure 1I-L,O,P). These live imaging studies confirmed that loss of *nudc* leads to accumulations of late endosomes and autophagosomes in axon terminals.

### Velocity of autophagosome and late endosome retrograde transport is reduced in *nudc* mutants

We reasoned that cargo accumulation in *nudc* mutant axon terminals could be due to inhibition of retrograde transport. To determine if transport was altered, we used live imaging of organelle transport in pLL axons in vivo [36]. First, we analyzed autophagosome retrograde transport using expression of RFP-tagged LC3. Neither the percent of autophagosomes moving in the retrograde direction (Figure 2A-C; Percent: WT – 34.14±2.6 v. *nudc* –35.19±2.9; ANOVA, *p*=0.7890) or the absolute number of autophagosomes in the axon were changed in *nudc* mutants (Figure 2D; WT – 30.14±3.39 v. *nudc–* 29.69±3.75; Wilcoxon). However, analyses of transport parameters revealed decreased retrograde distance and velocity in *nudc* mutants (Figure 2E,F). Histogram analyses of binned velocities demonstrated a clear shift in the population towards slower retrograde transport (Figure 2F’’).

**Figure 2.**
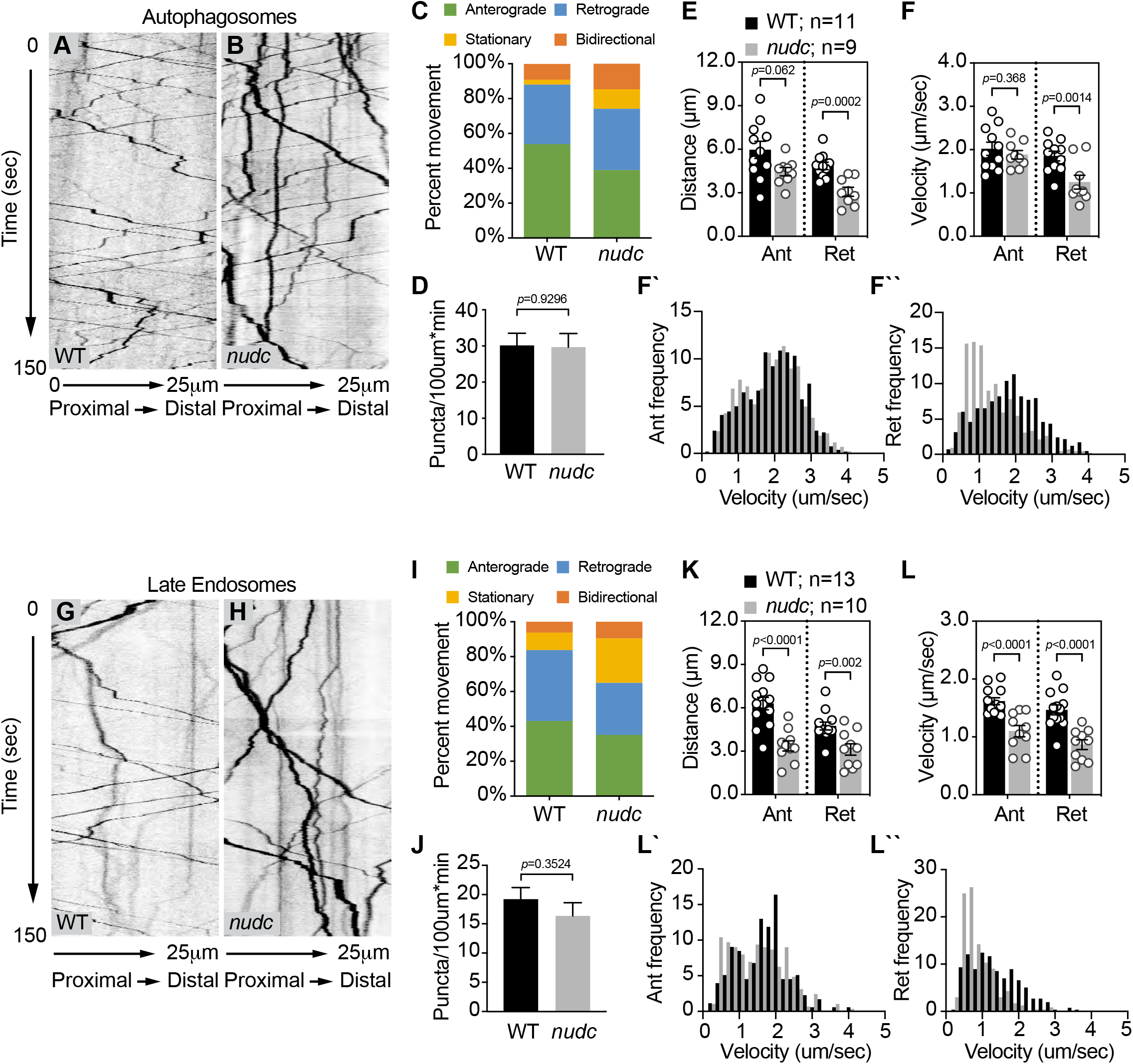
Decreased retrograde transport velocity of autophagosomes and late endosomes in *nudc* mutants. (A,B) Kymograph analysis of LC3-labeled autophagosome transport. (C) The autophagosome transport frequency in *nudc* mutants. Anterograde transport frequency decreases but the percent of autophagosomes moving in the retrograde direction is not changed (ANOVA; anterograde: *p*=0.0045; retrograde: *p*=0.7890). Stationary particle frequency increases (ANOVA; *p*=0.0023). (D) Total number of autophagosomes in the axon is unchanged (ANOVA). (E,F) Quantification of autophagosome transport distance and velocity show defects for both during retrograde transport (ANOVA). (F’,F’’) Histograms of binned transport velocities shows a clear shift to slower velocities for retrograde transport in *nudc* mutants. (G,H) Kymograph analysis of Rab7-labeled late endosome transport. (I) The percent of retrograde late endosome transport decreases in *nudc* mutants (ANOVA; p=0.0346), while stationary particle frequency increases (ANOVA; *p*=0.0031). (J) Total number of late endosomes in the axon is unchanged (ANOVA; *p*=0.3524). (K,L) Quantification of late endosome transport distance and velocity show defects in both the anterograde and retrograde directions (ANOVA). (L’,L’’) Binned histograms of transport velocities show a clear shift in retrograde transport towards slower speeds. Data are expressed as mean ± S.E.M. Sample sizes indicated on the graph.

Next, we analyzed late endosome transport using expression of RFP-tagged Rab7 in single pLL axons. Kymograph analysis demonstrated a slight but not significant decrease in retrograde transport frequency (Percent: WT – 40.65±3.11 v. *nudc* −29.98±3.55; Wilcoxon, *p*=0.0714) but no change in the total number of late endosomes in axons (WT – 19.20±1.98 v. *nudc–* 16.35±2.25; Wilcoxon); Figure 2G-J). Distance moved is decreased in both directions and a strong reduction of transport velocity, particularly in the retrograde direction, was also evident for late endosomes (Figure 2K,L). The clear shift of retrograde velocities towards slower speeds is apparent in the binned histogram (Figure 2L’’).

We were curious if these altered transport parameters were specific to these organelles or if similar deficits would be apparent for other cargos. Analyses of peroxisome and mitochondrial transport revealed no change in frequency of transport for these cargos; however, similar to the cargos analyzed above, a consistent reduction of mitochondrial velocity was observed (Figure S2). This was particularly obvious for retrograde transport with a shift of velocities towards slower speeds noted in the binned histogram analyses (Figure S2L’’). Finally, we asked if the movement of dynein-bound vesicles was generally impacted by loss of NudC. For this, we imaged transport of cargo labeled by Dync1li1V2 tagged with mRFP. Dync1li1V2 is the light intermediate chain variant shown to be present on autophagosomes and lysosomes being transported by dynein [37]. Consistent with our autophagosome and late endosome data, Dync1li1V2-labeled vesicles exhibited no change in retrograde transport frequency (WT – 29.48±3.78 v. *nudc–* 34.23±3.78; ANOVA, *p*=0.3839) or total numbers (WT – 22.41±2.50 v. *nudc–* 17.91±2.50; ANOVA) but did display decreased retrograde velocity (Figure S3). Together, the live imaging data support a model in which loss of NudC decreases retrograde transport processivity, contributing to ineffective retrograde transport and vesicle accumulation in axon terminals of *nudc* mutants.

### Axon terminal microtubule stability is enhanced in *nudc* mutants

One potential defect that could contribute to cargo accumulations and abnormal transport is a defect in the microtubule cytoskeleton. We began to address this by immunolabeling for total tubulin (Tubb3) and acetylated microtubules. Microtubule acetylation is tubulin modification associated with stabilized microtubules [1, 38]. We found no change in the mean fluorescence intensity (FI) for both Tubb3 and acetylated tubulin, indicating that total microtubule abundance in *nudc* axon terminals was unaltered. (Figure 3A-F). Though fluorescence intensity was unchanged, we did observe that *nudc* mutant axon terminals displayed a network of wrapped, acetylated microtubules (Figure 3E). This phenotype was not due to a change in axon terminal size as immunolabeling for acetylated tubulin in two other mutant strains with swollen axon terminals (*p150* and *actr10;* [33, 34]) did not show changes in acetylated microtubule structure (Figure 3G,H).

**Figure 3.**
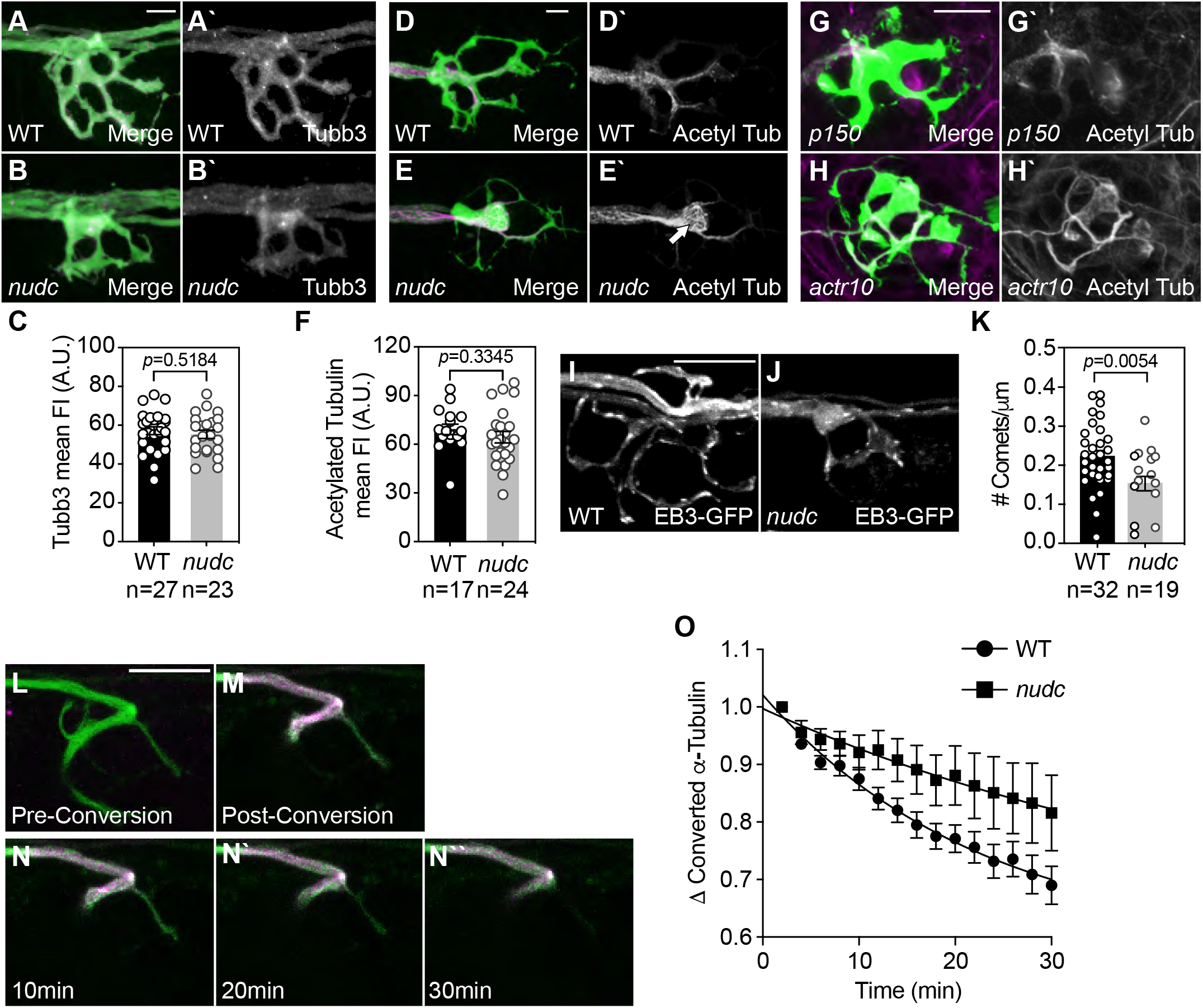
Microtubules in *nudc* axon terminals show signs of enhanced stabilization. (A,B) Staining of Tubb3 in wild type and *nudc* terminals. (C) Quantification of Tubb3 fluorescence intensity shows no difference in *nudc* terminals (ANOVA). (D,E) Labeling of acetylated tubulin in wild type and *nudc* terminals. White arrow in (E’) highlights the net-like organization of acetylated tubulin staining in mutants (inset). (F) Quantification of acetylated tubulin fluorescence intensity shows no difference in *nudc* terminals though organization is altered (ANOVA). (G,H) Acetylated tubulin labeling in axon terminals of two unrelated mutants, *p150* and *actr10*, show no change in microtubule structure despite changes in axon terminal structure. (I,J) Wild type (I) and *nudc* mutant (J) axon terminal showing EB3-GFP comets. (K) Axon terminal comet number is reduced in *nudc* mutants (ANOVA). (L-N) Dendra2-labeled α-tubulin in an axon terminal of the pLL. Prior to conversion, (L) no red fluorescence is detected (shown in magenta). Post-conversion, strong labeling of converted Dendra2-α-tubulin is observed (M). Converted Dendra2-α-tubulin decay over 30min post-conversion shown in N. (O) Decay curve of converted Dendra2-α-tubulin from time-lapse imaging post-conversion demonstrates a shift in the half-life in *nudc* mutants (mixed effects model; *p*=0.0033). Data are expressed as mean ± S.E.M. Scale bar = 10μm. Sample sizes indicated on graphs.

Dynein is well-known for its localization to microtubule plus ends [4, 39, 40]. This localization, which is at least partially controlled through the interaction of dynactin with microtubules and plus-end tracking proteins [39, 41], is thought to facilitate dynein-dynactin-cargo interaction in this region for the initiation of retrograde transport [2, 5, 6]. We reasoned that a change in microtubule plus end dynamics could contribute to the *nudc* mutant phenotype, so we next assessed microtubule stability directly in *nudc* mutant axon terminals. To determine if microtubule growth dynamics were decreased in the axon terminal, we analyzed microtubule plus ends labeled by the plus end binding protein EB3 tagged with GFP [42]. Due to the three-dimensional nature of the axon terminal, we could not effectively track EB3 comets to directly analyze microtubule dynamics. Instead, we analyzed comet number at a single timepoint. This analysis demonstrated a strong reduction in EB3 comet number with loss of NudC, suggesting decreased dynamic microtubule plus ends in the axon terminal (Figure 3I-K).

Though suggestive of increased microtubule stability, these single time-point analyses do not directly address persistence of microtubules in this region. To assess microtubule lifetime in the axon terminal, we turned to photoconversion. We expressed a-tubulin tagged with the photoconvertible protein Dendra2 in neurons [43]. Using 405nm light, we converted microtubules in the axon terminal branchpoint from green to red and then used live imaging to visualize red fluorescence decay, an indication of microtubule turnover (Figure 3L-N). After photoconversion, images were taken every two minutes for thirty minutes to ascertain the half-life of converted tubulin in this region. Similar to work in cultured hippocampal neurons, we observed a half-life of 16 minutes in wild type axon terminals [43]. In *nudc* mutants, the half-life of converted tubulin was more than double that of wild type siblings, roughly 36 minutes (Figure 3O; Rate constant: WT = 0.044; *nudc* = 0.019). These experiments indicate that microtubules in *nudc* mutant axon terminals are significantly more stable than in wild type siblings.

Changes in microtubule stability could underlie the transport defects observed in *nudc* mutants axons. Therefore, we asked if the increased microtubule stability was only in the axon terminal or also in the axon shaft as well. To address this, we again used EB3 tagged with GFP to analyze microtubule growth dynamics in the axon. There was no change in the frequency, distance, or velocity of microtubule growth in *nudc* mutants (Figure S4A-E). Microtubule structure could also be impacted independently of microtubule dynamics. To determine if microtubule organization was altered in *nudc* mutant axons, we labeled microtubule minus ends with the coiled-coiled domain of Patronin and analyzed their density [44]. This truncated form of Patronin has been shown to label minus ends without affecting minus end growth [44]. We expressed PatroninCC in pLL axons and measured PatroninCC punctal density in the cell body, axon, and axon terminal. This revealed no difference between WT and *nudc* mutants (Figure S4F-J). These data suggest that though axon terminal microtubules are stabilized in *nudc* mutants, microtubule dynamics and structure in the axon shaft are unaffected and do not contribute to transport phenotypes.

### Loss of Lis1 in axon terminals underlies *nudc* phenotypes

The best characterized function for NudC is in regulation of Lis1 protein stability through NudC’s chaperone function [24, 30]. Lis1 promotes the formation of active dynein complexes for retrograde transport through promoting an “open” rather than autoinhibited conformation [12, 13]. This promotes double dynein dimer binding to cargo [10, 11] which can enhance retrograde transport velocities [45]. Retrograde transport velocities are reduced in *nudc* mutants, so we hypothesized that decreased Lis1 stability in the absence of NudC contributed to the cargo localization and transport abnormalities in *nudc* mutants. We investigated the stability and localization of Lis1 in *nudc* mutant neurons. First, we assessed Lis1 protein from whole larval extracts. Western blot analysis revealed an almost complete loss of Lis1 protein in *nudc* mutants (Figure 4A,B). Next, we analyzed Lis1 levels in neurons of intact animals. We could not optimize parameters for an anti-Lis1 antibody for whole mount immunolabeling. Therefore, to address this question, we turned to overexpression. We expressed Lis1 tagged with mRFP in neurons and analyzed fluorescent intensity. This analysis revealed no significant reduction of Lis1 in the cell body but a strong reduction of Lis1-mRFP protein in axon terminals (Figure 4C-E). Together, these data suggest a decrease in Lis1 protein in *nudc* mutants, with a more notable loss in the distal axon terminal.

**Figure 4.**
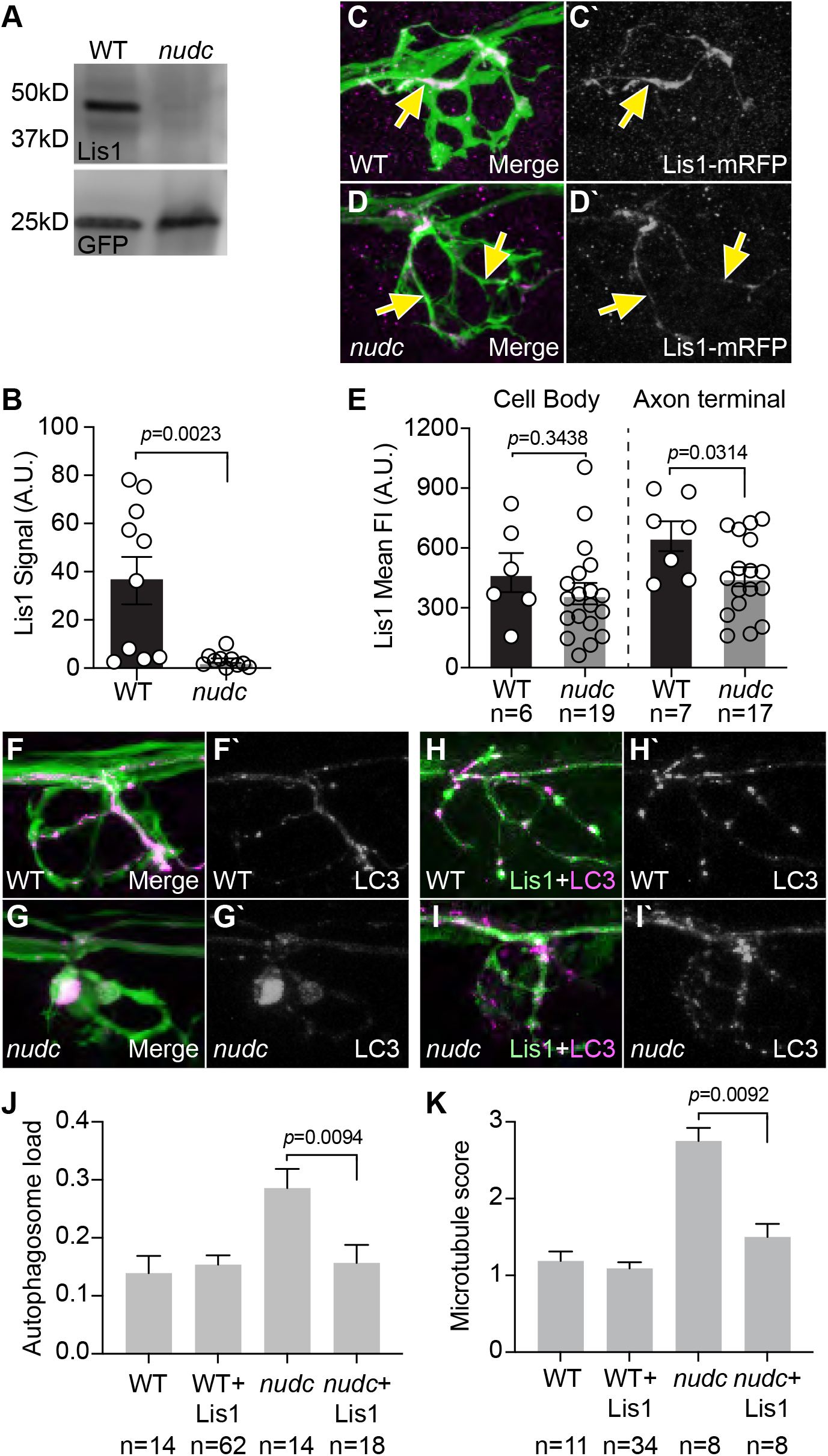
Lis1 functions downstream of NudC in regulation of cargo transport and microtubule stability. (A) Lis1 western demonstrates loss of Lis1 protein in *nudc* mutants at 4 dpf. GFP from *TgBAC*(*neurod:egfp*)*^nl1^* transgene serves as an internal control. (B) Quantification of signal intensity relative to blot background confirms loss of Lis1 in *nudc* mutants (n=10 independent blots; ANOVA). (C,D) Expression of Lis1-mRFP in individual neurons (magenta in merge, white on right). Arrows point to axons expressing Lis1. (E) Quantification of Lis1 mean fluorescence intensity in the cell body and axon terminal of *nudc* mutants and wild type siblings demonstrates a strong loss of Lis1 in mutant axon terminals (ANOVA). (F-I) Expression of Lis1-eGFP can suppress autophagosome accumulation in *nudc* mutants. (F,G) Expression of mRFP-LC3 (magenta in merge, white on right) in single axons of a wild type (F) and *nudc* mutant (G) shows accumulation in *nudc* mutants. Both carry the *TgBAC*(*neurod:egfp*)^*nl1*^ transgene to label neurons with cytosolic GFP (H,I) Co-expression of Lis1-eGFP and mRFP-LC3 (magenta in merge, white on right) in *nudc* mutant axons can suppress autophagosome accumulation. (J) Quantification of autophagosome load (autophagosome area/axon area) demonstrates suppression of accumulation by expression of Lis1-eGFP in *nudc* mutants (ANOVA Tukey HSD post-hoc contrasts). (K) Expression of Lis1-eGFP can suppress netting of acetylated microtubules in *nudc* mutants (scale: 1 - wild type microtubules; 2 – mild microtubule looping; 3 – presence of microtubule nets; Wilcoxon each pair). Data are expressed as mean ± S.E.M. Scale bar = 10μm. Sample sizes indicated on graphs.

Next, we wanted to determine whether loss of Lis1 was responsible for the cargo accumulation phenotypes we observed in *nudc* mutants. To address this, we attempted to rescue autophagosome localization with exogenous Lis1 expression. We overexpressed Lis1 tagged with eGFP in neurons and analyzed autophagosome density in the axon terminal using mRFP-tagged LC3. Overexpression of Lis1 suppressed autophagosome accumulation in *nudc* mutant axon terminals (Figure 4F-J). Together, this data suggests that loss of Lis1 in *nudc* mutants leads to the accumulations of cargo in the axon terminal.

We then asked if loss of Lis1 caused the microtubule stabilization phenotype in *nudc* mutants. We expressed Lis1 (tagged with eGFP) and immunolabeled *nudc* mutants and wild type siblings with anti-acetylated tubulin antibodies to assess the microtubule cytoskeleton in axon terminal. Microtubule structure was imaged and quantified on a rank order scale by two independent blinded reviewers (1 - normal microtubules; 2 – mild microtubule looping; 3 – presence of microtubule nets). This analysis revealed that exogenous Lis1 also suppressed the formation of stabilized microtubule nets in *nudc* mutant axon terminals (Figure 4K). Together, our data suggest that NudC-mediated Lis1 stability in the axon terminal is crucial for regulating both efficient retrograde cargo transport and microtubule stability.

### Dynein and dynactin components accumulate in *nudc* mutant axon terminals

In addition to regulating Lis1, previous biochemical studies have implicated NudC in the regulation of kinesin-dependent dynein anterograde transport [46]. Loss of dynein motor components from the axon terminal could contribute to cargo accumulation independent of Lis1 deficits. To address this, we analyzed dynein localization in the axon terminal using immunolabeling and live imaging. First, we labeled axon terminals with antibodies specific for dynein heavy chain and p150, a core component of dynactin. Fluorescence intensity analysis demonstrated an increase of both components of the retrograde motor in *nudc* axon terminals (Figure 5A-E). We next analyzed the localization of dynein-labeled vesicles in vivo using live imaging. We expressed dynein light intermediate chain 1 variant 2 (Dync1li1V2) tagged with mRFP in pLL neurons to assess the localization of vesicles labeled with this dynein subunit. This also revealed an increase in this dynein component in axon terminals (Figure 5F-H). Together, these assays of endogenous and overexpressed dynein-dynactin components show that anterograde movement of dynein-dynactin proteins is intact. Furthermore, it reveals that these motor components accumulate in axon terminals of *nudc* mutants.

**Figure 5.**
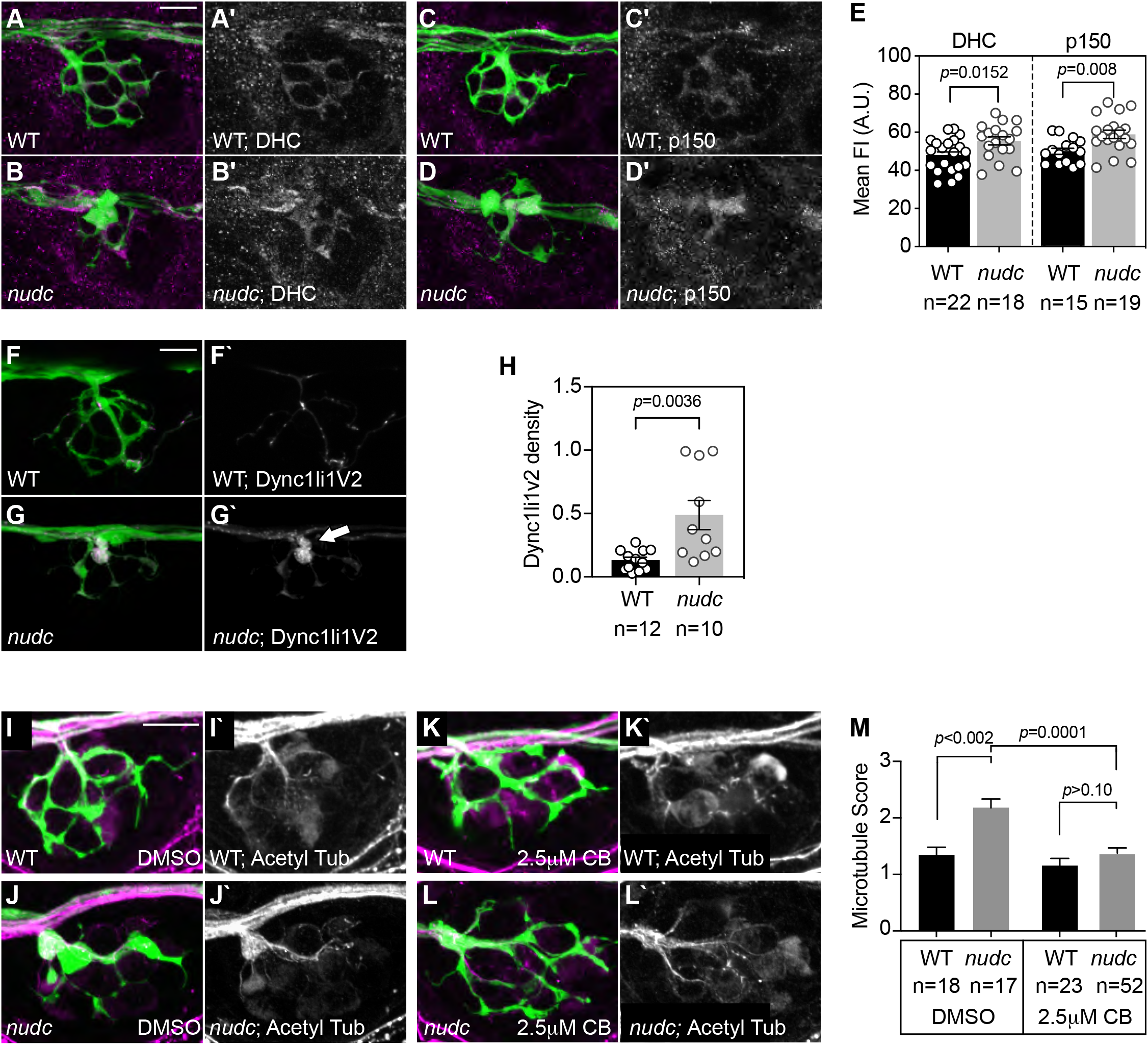
Pharmacological inhibition of dynein suppresses microtubule stabilization in *nudc* mutant axon terminals. (A-D) Dynein heavy chain (DHC) and p150 (dynactin) accumulate in *nudc* mutant axon terminals. (E) Quantification of mean fluorescence intensity (normalized to background) for DHC and p150 (ANOVA). (F,G) mRFP-Dync1li1V2 expression in individual axons of wild type and *nudc* mutants demonstrates accumulation in mutant axon terminals (arrow). (H) Quantification of mRFP-Dync1li1V2 density (Dync1li1V2 area/axon terminal area) demonstrates accumulation of this dynein light intermediate chain in *nudc* mutant axon terminals (ANOVA). (I-L) Wild type (I) and *nudc* mutant (J) axon terminals after 24hr 0.1% DMSO treatment. *nudc* mutants show large axon branchpoint swellings and acetylated tubulin netting in those swellings. (K,L) Treatment with 2.5μM ciliobrevin (CB) suppresses axonal swellings and microtubule netting in *nudc* mutants. (M) Average qualitative score from two independent, blinded analyses (scale: 1 - wild type microtubules; 2 – mild microtubule looping; 3 – presence of microtubule nets; Wilcoxon each pair). Data are expressed as mean ± S.E.M. Scale bar = 10μm. Sample sizes indicated on graphs.

### Dynein-dependent regulation of microtubule stability in the axon terminal

Dynein, in the presence of ATP, has been shown to stabilize microtubules in vitro and loss of dynein leads to loss of microtubule stability in C. elegans dendrites in vivo [14, 15]. We hypothesized that the accumulation of dynein in the axon terminal of *nudc* mutants could underlie the increase in microtubule stability. To test this, we inhibited dynein activity with a low dose of ciliobrevin (used previously in zebrafish; [47]) and assessed microtubule structure with immunolabeling for acetylated tubulin. Microtubule structure was quantified blind by two independent reviewers on a rank order scale (1 - normal; 2 – mild microtubule looping; 3 – presence of microtubule nets) and larvae were genotyped post-analysis. Strikingly, ciliobrevin treatment could suppress axon terminal swellings and formation of stabilized microtubule nets in the axon terminal of *nudc* mutants (Figure 5I-M). This indicates that increased dynein activity in the axon terminal causes the increased microtubule stability phenotype in *nudc* mutants.

## Discussion

Investigation of dynein function in the axon terminal has largely focused on the regulation of motor-cargo attachment and initiation of retrograde transport from the terminal towards the cell body. This process is critical for neuronal health and function. Our work has re-emphasized a second role for dynein in this compartment, regulation of the microtubule cytoskeleton. In this study, we show that NudC is critical for Lis1 protein stability. Similar to work in Aspergillus [23, 24], loss of NudC causes a strong reduction in Lis1 protein which is particularly evident in the distal axon terminal compartment. Based on the recent advances in our understanding of Lis1 function [10–13] and the presented work, loss of Lis1 in *nudc* mutants results in decreased dynein-cargo interaction and initiation of retrograde transport from the axon terminal. Based on this premise, our data support a model in which NudC-dependent Lis1 stability is essential for dynein-cargo interaction for retrograde transport from the axon terminal. Disrupting this process leads to accumulation of commonly transported cargo and the dynein motor. We propose that it is this accumulation of the dynein-dynactin complex in the axon terminal which, when its activity is left unchecked, enhances microtubule stability in this neuronal compartment.

### Regulation of the axon terminal microtubule cytoskeleton by NudC

The axonal cytoskeleton is a subject of intensive research, particularly during axon extension. During axon outgrowth, the growth cone has an array of actin and microtubule dynamics that allow environmental exploration to search for and innervate distant targets. In the growth cone, active actin retrograde flow and microtubule instability allow the constant remodeling of this regions for rapid response to extracellular cues (reviewed in [1, 48]). Though less well studied, the terminal of a mature axon also requires tight regulation of actin and microtubule structure and dynamics. Microtubule plus ends reside in the axon terminal and contribute substantially to dynein recruitment and dynein-cargo interaction for the initiation of retrograde transport [2, 5, 6]. These plus ends are highly dynamic relative to the rest of the axon, with shorter half-lives and markers of unstable microtubules including tyrosination [1, 38]. Further emphasizing the importance of regulating microtubule dynamics in the neuron, increasing microtubule stability is associated with neurodegeneration. Treatment with the microtubule stabilizing drug taxol, a common pharmacological therapeutic for cancer, results in axonal degeneration and peripheral neuropathy in patients. Additionally, mutations in the microtubule severing protein Spastin cause the degenerative disease hereditary spastic paraplegia (reviewed in [49]). Despite the importance of microtubule dynamics in the axon terminal, we know little about the roles they play in the maintenance of a functional neural circuit.

Our work emphasizes a role for dynein in the control of microtubule stability in the axon terminal. In *nudc* mutants, dynein and p150 accumulate in axon terminals and microtubules in the axon terminal show signs of enhanced stabilization. These signs include: 1) microtubule acetylation, a marker more stable microtubules in neurons [1, 38]; 2) longer half-life of tubulin in the region based on photoconversion data; and 3) fewer EB3-GFP comets which are a label for dynamic microtubule plus ends. Decreasing dynein activity using ciliobrevin treatment can suppress microtubule stabilization, implicating excessive dynein activity in this phenotype. This work is in-line with previous work in *C. elegans* dendrites and in vitro which has demonstrated that dynein can stabilize microtubules. In *C. elegans*, expression of mutant forms of dynein associated with neurological disease leads to decreased microtubule stability [15]. In vitro, isolated microtubules undergo catastrophe when they contact a barrier. Including dynein and ATP in this system prevents microtubule depolymerization upon barrier contact [14]. Similar to the growth cone [21], microtubule plus ends are likely in constant contact with barriers such as the cellular cortex and actin cytoskeleton in the mature axon terminal. In addition to dynein, the p150 subunit of dynactin has also been strongly implicated in microtubule stabilization in yeast, in vitro, and in mammalian neurons [17, 50–53]. Finally, NudC has independently been implicated in microtubule regulation during cell division in mammalian cells in culture and *C. elegans*, though whether this is due to regulation of dynein is unknown [32]. Together, these studies and our data suggest that it is the increase in local dynein-dynactin activity that stabilizes microtubules in *nudc* mutant axon terminals, allowing them to resist depolymerizing forces, such as contact with barriers. It is likely that this function relies on microtubule-cell cortex interaction; however, proteins involved in this interaction are unknown.

### Dynein-dynactin function in microtubule regulation

Cytoplasmic dynein has previously been implicated in the regulation of microtubule stability in neurons and nonneuronal cells. However, a clear picture of the role of this motor in microtubule regulation is still lacking. This is evidenced by the conflicting data on dynein’s action on microtubules. In yeast, fungus, and non-neuronal cells, dynein decreases microtubule stability, particularly when dynein is attached to the cellular cortex and acting on microtubule plus ends [16–19]. In yeast, when plus end-bound dynein is attached to the cell cortex, the motor increases microtubule dynamics through promoting microtubule catastrophe [17, 19]. The resulting microtubule shrinkage is capable of creating a pulling force which facilitates centralized microtubule aster positioning in vitro [19] and likely contributes to microtubule-dependent spindle and nuclear positioning during cell division in yeast and *C. elegans* [54, 55]. Similarly, in *Aspergillus*, loss of dynein function causes increased microtubule growth into the hyphal tip and decreased microtubule catastrophe and shrinkage rate [18]. These results are suggestive of a role for dynein in decreasing microtubule stability upon interaction between the microtubule plus end and a barrier.

Conversely, dynein-mediated microtubule tethering to the cell cortex has also been implicated in microtubule stabilization. In vitro, though initially increasing microtubule catastrophe frequency, once the microtubule is pulled taught, dynein-mediated microtubule tethering to a barrier actually decreases microtubule shrinkage rate [19]. There is also evidence that dynein on microtubule plus ends tethers microtubules to the cell cortex for stabilization in mammalian cells. In cultured epithelial cells, dynein has been shown to interact with β-catenin to help stabilize microtubules at the cell membrane [56]. Finally, work in vitro on mammalian dynein has also substantiated a role for the dynein-dynactin complex in increasing microtubule stability. Upon encountering a barrier, single microtubules will undergo catastrophe. However, when dynein and ATP were added, microtubules could withstand force induced through interaction with the barrier [14]. Together, these data argue that dynein can also serve a stabilizing role for microtubules.

In neurons, the existing evidence suggests that dynein primarily serves a stabilizing role for microtubules, particularly in axons and dendrites. In dendrites of *C. elegans* neurons, expression of loss of function dynein heavy chain mutants decreases microtubule stability in dendrites [15]. In axonal growth cones of cultured mammalian neurons, inhibition of dynein results in failed microtubule growth into the growth cone. In the growth cone microtubules cannot resist the rapid retrograde actin flow present in this region without dynein [21]. Furthermore, dynein interaction with the neural cell adhesion molecule NCAM180 has been found to tether dynamic microtubule ends at synapses and promote synaptic stability [60]. Similar to work in vitro and in yeast, the p150 subunit of dynactin has also been shown to promote microtubule stability in neurons as well [17, 51, 52]. This function for p150 may also be cell-type specific as knockdown of p150 has no effect on microtubule dynamics in cultured epithelial cells but has a strong destabilizing effect on microtubules in cultured neurons [52]. This suggests that there are context dependent controls in place which push dynein-dynactin into either a stabilizing or destabilizing role. These likely involve adaptor and modulatory proteins, localization of dynein to the lateral surface or plus end of the microtubule, and the mechanical forces at play on the microtubule.

### Lis1 and neuronal phenotypes in *nudc* mutants

NudC has been implicated in Lis1 protein stability previously in vitro and in *Aspergillus* [23, 24, 29, 63]. Our data support this function for NudC. In whole larval extracts, Lis1 protein levels are severely reduced. Using overexpression of tagged Lis1, we show that Lis1 protein is reduced throughout the neuron; however, there is a more severe reduction in the distal axon terminal. This is likely due to the long distance from the cell body to the axon terminal in this system. Unless Lis1 protein is produced locally in the axon terminal, it must be synthesized and transported from the cell body to the axon terminal. Indeed this appears to be the case as recent RNAseq-based approaches have shown a biased presence of Lis1 mRNA in neuronal cell bodies rather than the neuropil [64]. This further supports the need to preserve Lis1 protein stability particularly once it arrives in the axon terminal for function.

NudC-dependent Lis1 stability is critical for dynein regulation. Lis1 has recently been shown to be critical for dynein-cargo interaction for processive retrograde transport through promotion of the open dynein conformation and subsequent dual dynein dimer interaction with dynactin and cargo [10–13]. In addition to cargo transport, Lis1 has also been implicated in the regulation of microtubule stability. Inhibition of Lis1 in neurons results in failed microtubule stabilization and penetration of the growth cone necessary for growth cone advance [21]. This could be through direct interaction of Lis1 with microtubule plus end proteins including CLIP-170 [20, 22] or through modulation of dynein-dynactin function. Our data suggest that Lis1-dependent microtubule modulation in the axon terminal is indirect through Lis1’s effect on dynein. In our experiments, microtubule dynamics in *nudc* mutants can be rescued by exogenous expression of Lis1 and by suppressing dynein activity. Together, this suggests that in the axon terminal the bulk of Lis1 function is in regulating dynein activity which has subsequent effects on cargo localization and microtubule stability.

The dramatic defects on microtubules in *nudc* mutant axon terminals suggests that the NudC-Lis1-dynein cascade must be spatially and temporally controlled to maintain the structure and function of the axon terminal. How does NudC tightly regulate Lis1-mediated dynein activation? Some potential hints at NudC control come from studies on the function of this protein during cell division. In *C. elegans* and cultured mammalian cells, NudC is essential for microtubule stability at the midzone which is essential for cell division. NudC’s function during cell division relies on interaction with Plk1 (Polo-like Kinase 1) [32]. Plk1 phosphorylates NudC leading to translocation of the NudC-Plk1 complex to the kinetochore. Loss of NudC phosphorylation by Plk1 prevents this translocation and proper cell division [61, 62]. Therefore, phosphorylation of NudC is poised to function as a regulator of localized NudC function. Whether NudC phosphorylation impacts axonal phenotypes in mature neurons has not been explored but is a clear direction of interest for precise regulation of dynein function in neuronal compartments.

## Acknowledgements

Funding for this work was provided by the Provost and the Office of the Vice Chancellor for Research and Graduate Education at the University of Wisconsin-Madison with funding from the Wisconsin Alumni Research Foundation (to C.M.D.), the NINDS (NS111419 to A.V.N), and the NIH Intramural Research Program Grants from the NICHD (1ZIAHD008964-02 to C.M.D.) and the NIDCD (DC000081 to R.S.P. and Y.-X. W.). We want to thank A. Mandal, K. Klier, S. Wisner, H-T. C. Wong, C. Stein, A. Lang, and D. Mai for their thoughtful comments on this work. We would like to also acknowledge E.P. Louise for his suggestion to try the dynein inhibition experiment. PatroninCC and LC3 constructs were acquired from Addgene. The authors declare no competing financial interests.

## Author Contributions

DK, KP, and MC designed, performed, and analyzed experiments. RSP and YXW designed and performed electron microscopy experiments. AVN designed and facilitated the forward genetic screen to identify the *nudc* mutant. CMD supervised the study, designed, performed, and analyzed experiments. CMD wrote the paper with input from DK, KP, AVN, and RSP.

## STAR Methods

### Resource availability

Further information and requests for resources and reagents should be directed to Dr. Katie Drerup. All resources will be made available upon request.

### Experimental Models

#### Zebrafish husbandry

All zebrafish (*Danio rerio*) work was done in accordance with the University of Wisconsin-Madison IACUC guidelines. The *TgBAC*(*neurod:egfp*)*^nl1^* transgenic line was utilized for many of the studies. Adult animals were kept at 28°C and spawned according to established protocols [68]. Embryos and larvae were kept in embryo media at 28°C and developmentally staged using established methods [69].

### Method Details

#### Generation and genotyping of the nudc^nl21^ mutant strain

N-ethyl-N-nitrosourea (ENU) mutagenesis was done using the *TgBAC*(*neurod:egfp*)*^nl1^* transgenic as previously described [31]. The *nudc^nl21^* mutant was identified in a three generation, recessive mutant screen targeted to find mediators of retrograde cargo transport through their axon terminal swelling phenotype. The mutation in the *nudc* strain was identified as a T to A mutation in the splice donor site of intron 8 using RNAmapper [70]. This mutation was confirmed by sequencing *nudc* cDNA from mutant larvae. A combination of genotyping primers can be used to genotype this locus by PCR followed by Xmn1 digest. The *nudc^nl21^* mutation inserts an Xmn1 cut site. Reverse transcription of wildtype mRNA was done using the Superscript IV kit (Invitrogen).

#### Transient transgenesis

For analyses of cargo localization in single neurons, transient transgenesis was utilized. Plasmid DNA encoding the *5kbneurod* (pLL neurons; [71] promotor driving constructs of interest were derived using Gateway technology [72]. For expression, 3-13pg of plasmid DNA was microinjected into zebrafish zygotes as previously described [31, 36]. For analysis, larvae expressing the construct of interest in a subset of pLL neuron cell bodies were selected using a Zeiss AxioZoom fluorescent dissecting microscope. Larvae were anesthetized in 0.02% tricaine and mounted individually in 1.5% low melt agarose in embryo media and imaged with a Zeiss LSM800 or Olympus FV3000 confocal microscope with a 63X/NA1.2 water immersion or 40X/NA1.2 silicone immersion objective respectively. RFP-Rab7, Mito-TagRFP, EB3-GFP, PST1-RFP (peroxisomes), and RFP-Dync1li1v2 constructs for transient transgenesis have been used previously [31, 34]. Human LC3 was subcloned for expression from Addgene clone 24920 [73]. PatroninCC was subcloned from the Addgene clone 59053 [44]. *nudc and lis1b* cDNAs were subcloned from zebrafish cDNA and confirmed by sequencing.

#### Immunohistochemistry and western blot

Whole mount immunohistochemistry on zebrafish larvae was done as previously described [31]. Briefly, zebrafish larvae at 4 dpf were fixed in 4% paraformaldehyde/0.1% triton overnight at 4°. After washes in 1X PBS/0.1% triton, larvae were washed in RNAse/DNAse-free water overnight at room temperature before blocking in 5% goat serum. Larvae were then incubated in primary antibody in blocking solution overnight at 4°, washed in 1X PBS/0.1% triton, incubated in Alexafluor secondary antibodies overnight at 4°, washed again in 1X PBS/0.1% triton, and sunk in 60% glycerol. Immunolabeled larvae were imaged on a LSM800 Zeiss or FV3000 confocal microscope with a 63X/NA1.4 or 60X/NA1.4 objective respectively. A 1024X1024 frame size was used and the image was generated by 2X averaging of the frame.

Western blot was done as previously described [34]. Protein lysates were run on a 4-20% SDS-PAGE denaturing gel before being transferred to PVDF membrane. Membranes were blocked in 5% Omni-block in 1X PBS/0.1% Tween 1-4 hours before being incubated in primary antibody overnight in block at 4°. Membranes were washed in 1X PBS/0.1% tween 3 times for 5 minutes before being incubated in secondary antibody (Goat anti-rabbit (Jackson Immuno Research): 1:10000) or 90 minutes at room temperature. Membranes were washed again 3 times for 5 minutes and developed using WesternSure Chemiluminescent Substrate on a C-DiGit blot scanner (Licor). Quantification of western blots using densitometry was done in Image Studio. Mean intensity was determined for each band and normalized to the background intensity across the blot.

#### Electron microscopy

Larvae (4 dpf) were prepared for transmission electron microscope (TEM) study as described previously [74, 75]. Briefly, larvae were fixed in 4% paraformaldehyde plus 2% glutaraldehyde (Electron Microscopy Sciences (EMS)) in phosphate buffer, then washed in cacodylate buffer, fixed in 2% glutaraldehyde, and washed again in cacodylate buffer. Then they were placed in 1% osmium tetroxide in cacodylate buffer, washed again in cacodylate buffer, and then dehydrated in an ethanol series with 1% uranyl acetate added to the 50% ethanol. Larvae then were placed in propylene oxide (PO), and then in an epon/PO mix and finally in pure epon, and samples were embedded and then hardened in an oven at 64°. Thin sections (60 nm) were placed on single-slot, formvar/carbon coated nickel grids (EMS), stained with uranyl acetate and lead citrate, and examined in a JEOL JEM2100 TEM.

#### Quantification of immunofluorescence

For quantification of immunofluorescence, whole mount, immunolabeled larvae were imaged using an LSM800, 63X/NA1.4 objective (Zeiss) or FV3000, 60X/NA1.4 objective (Olympus) confocal microscope. Laser power and detector gain were kept consistent for all images acquired for quantification of immunofluorescence. Quantification was done in ImageJ. For quantification, images from the 488 nm (neuronal) and 568 nm (antibody label of interest) were projected into 2D images using the Sum Stacks *z*-projection feature. The neuronal area was selected in the axon terminal by thresholding the 488 nm z-stack and using the ImageJ image subtraction plugin to remove non-neuronal signal from the 568nm channel. The mean fluorescence intensity of the 568nm channel was then quantified in a sum projection (ImageJ) and normalized to background.

#### In vivo analysis of cargo localization and axonal transport in the zebrafish pLL

The localization and transport of various cargos was done as previously described [36]. Fluorescently labeled cargos were expressed mosaically in neurons as described above using transient transgenesis. Larvae were mounted in 1.5% low melt agarose in embryo media on a coverslip after light anesthesia in 0.02% tricaine. For analysis of punctal volume, axon terminals expressing the tagged cargo of interest were imaged using an LSM800 63X/NA1.2 water immersion objective (Zeiss) or FV3000 40X/NA1.2 silicone immersion objective (Olympus). Laser power and detector gain were changed to identify puncta clearly based on expression level. For quantification, the area around the expressing terminal was cropped and the 488 nm (neuronal) and 568 nm (cargo) were split. Total punctal area and axon area were measured using built in ImageJ plugins. Within larval averages were used for analysis. Larvae were individually genotyped after imaging.

For analysis of cargo transport, imaging in a single plane was done on larvae mounted as described above on a LSM800 63X/NA1.2 water immersion objective (Zeiss) or FV3000 40X/NA1.2 silicone immersion objective (Olympus). A 25-75 μm length of axon was imaged for 500 frames with a 300 millisecond frame interval. Axonal transport was analyzed using kymograph analysis in Metamorph (Molecular Devices). Distance and time of individual bouts of transport were quantified from the kymographs. Number of puncta moving in the anterograde or retrograde direction as well as the number of bidirectional puncta were quantified using kymographs as well. Stationary puncta number was quantified by visual analysis of the transport imaging sessions. Anterograde or retrograde transport bouts were defined as traces representing puncta that only moved in one direction over the course of the imaging session.

For analysis of EB3-labeled microtubule growth dynamics, larvae were mounted and imaged as described above for axonal transport. The only modifications to the protocol were that images were taken every 15 seconds allowing *z*-stacks with an optimal *z*-step to be taken through the axonal region. For analysis of EB3 comets in the axon terminals, a *z*-stack was taken through the axon terminal and comet number was calculated manually from a projected optimal *z*-stack.

#### Ciliobrevin treatment and scoring of axon terminal morphology and acetylated microtubule structure

At 3 dpf, larvae from a *nudc* heterozygous cross were sorted into groups of 50. These larvae expressed the *TgBAC*(*neurod:egfp*)*^nl1^* transgene to visualize pLL axons. Each group was housed in a well of a 6-well plate. At 3.5 dpf, the embryo media in the well was replaced with embryo media with 0.1% DMSO or 2.5 μM ciliobrevin/0.1% DMSO. At 4.5 dpf, larvae were immunolabeled prior to imaging axon terminals on a Zeiss LSM800 confocal microscope with a 63X/NA1.4 oil immersion objective. Projected z-stacks were compiled for analysis by two blinded reviewers. Reviewers scored microtubule structure in axon terminals. 1 - wild type microtubules; 2 – mild microtubule looping; 3 – presence of microtubule nets. After imaging and analysis, larvae were individually genotyped for the *nudc* mutation.

### Quantification and Statistical analysis

For all experiments, all statistical analyses were done using JMP15. For parametric analyses, ANOVAs were used with Tukey HSD post-hoc contrasts for multiple pairwise comparisons within a dataset. For non-parametric datasets, comparisons were done using Wilcoxon/Kruskal-Wallis analyses with corrected tests for each pair for multiple comparisons.

## KEY RESOURCES TABLE

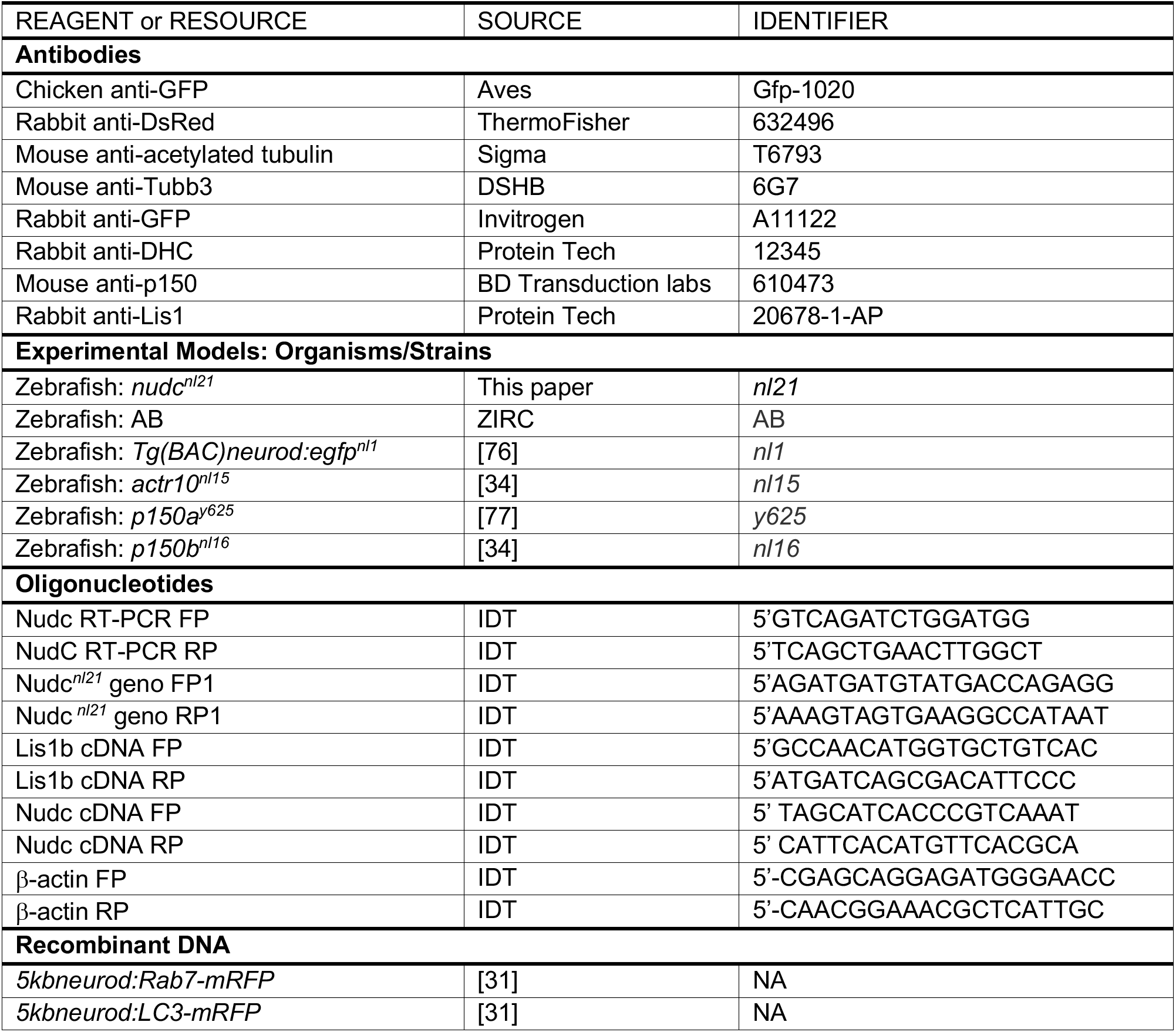
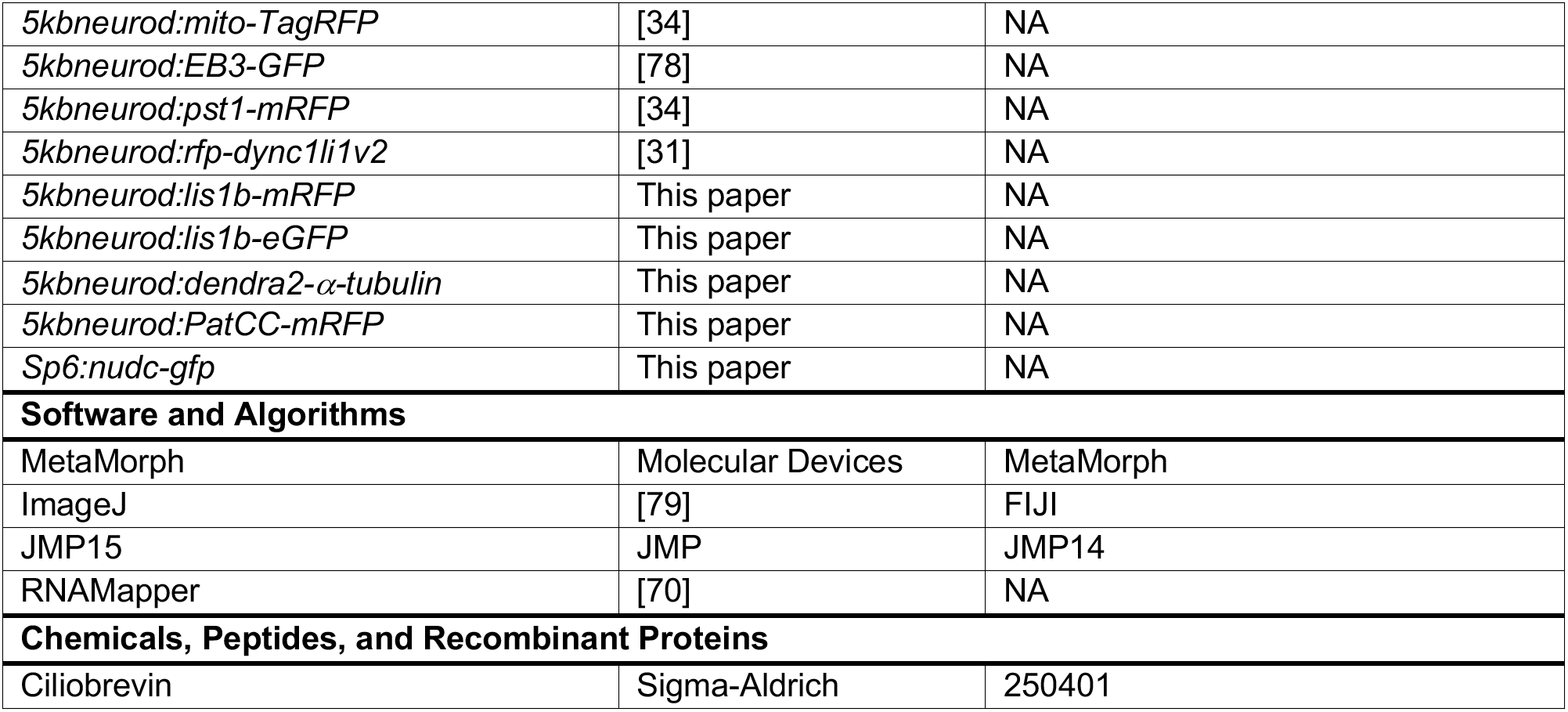

## References

1. Baas, P.W., Rao, A.N., Matamoros, A.J., and Leo, L. (2016). Stability properties of neuronal microtubules. Cytoskeleton (Hoboken) 73, 442–460.

2. Moughamian, A.J., Osborn, G.E., Lazarus, J.E., Maday, S., and Holzbaur, E.L. (2013). Ordered recruitment of dynactin to the microtubule plus-end is required for efficient initiation of retrograde axonal transport. J Neurosci 33, 13190–13203.

3. Splinter, D., Razafsky, D.S., Schlager, M.A., Serra-Marques, A., Grigoriev, I., Demmers, J., Keijzer, N., Jiang, K., Poser, I., Hyman, A.A., et al. (2012). BICD2, dynactin, and LIS1 cooperate in regulating dynein recruitment to cellular structures. Mol Biol Cell 23, 4226–4241.

4. Vaughan, K.T., Tynan, S.H., Faulkner, N.E., Echeverri, C.J., and Vallee, R.B. (1999). Colocalization of cytoplasmic dynein with dynactin and CLIP-170 at microtubule distal ends. J Cell Sci 112 (Pt 10), 1437–1447.

5. Lloyd, T.E., Machamer, J., O’Hara, K., Kim, J.H., Collins, S.E., Wong, M.Y., Sahin, B., Imlach, W., Yang, Y., Levitan, E.S., et al. (2012). The p150(Glued) CAP-Gly domain regulates initiation of retrograde transport at synaptic termini. Neuron 74, 344–360.

6. Moughamian, A.J., and Holzbaur, E.L. (2012). Dynactin is required for transport initiation from the distal axon. Neuron 74, 331–343.

7. Olenick, M.A., and Holzbaur, E.L.F. (2019). Dynein activators and adaptors at a glance. J Cell Sci 132.

8. Reck-Peterson, S.L., Redwine, W.B., Vale, R.D., and Carter, A.P. (2018). The cytoplasmic dynein transport machinery and its many cargoes. Nat Rev Mol Cell Biol 19, 382–398.

9. Urnavicius, L., Lau, C.K., Elshenawy, M.M., Morales-Rios, E., Motz, C., Yildiz, A., and Carter, A.P. (2018). Cryo-EM shows how dynactin recruits two dyneins for faster movement. Nature 554, 202–206.

10. Elshenawy, M.M., Kusakci, E., Volz, S., Baumbach, J., Bullock, S.L., and Yildiz, A. (2020). Lis1 activates dynein motility by modulating its pairing with dynactin. Nat Cell Biol 22, 570–578.

11. Htet, Z.M., Gillies, J.P., Baker, R.W., Leschziner, A.E., DeSantis, M.E., and Reck-Peterson, S.L. (2020). LIS1 promotes the formation of activated cytoplasmic dynein-1 complexes. Nat Cell Biol 22, 518–525.

12. Qiu, R., Zhang, J., and Xiang, X. (2019). LIS1 regulates cargo-adapter-mediated activation of dynein by overcoming its autoinhibition in vivo. J Cell Biol 218, 3630–3646.

13. Marzo, M.G., Griswold, J.M., and Markus, S.M. (2020). Pac1/LIS1 stabilizes an uninhibited conformation of dynein to coordinate its localization and activity. Nat Cell Biol 22, 559–569.

14. Hendricks, A.G., Lazarus, J.E., Perlson, E., Gardner, M.K., Odde, D.J., Goldman, Y.E., and Holzbaur, E.L. (2012). Dynein tethers and stabilizes dynamic microtubule plus ends. Curr Biol 22, 632–637.

15. Yogev, S., Maeder, C.I., Cooper, R., Horowitz, M., Hendricks, A.G., and Shen, K. (2017). Local inhibition of microtubule dynamics by dynein is required for neuronal cargo distribution. Nat Commun 8, 15063.

16. Carminati, J.L., and Stearns, T. (1997). Microtubules orient the mitotic spindle in yeast through dynein-dependent interactions with the cell cortex. J Cell Biol 138, 629–641.

17. Estrem, C., Fees, C.P., and Moore, J.K. (2017). Dynein is regulated by the stability of its microtubule track. J Cell Biol 216, 2047–2058.

18. Han, G., Liu, B., Zhang, J., Zuo, W., Morris, N.R., and Xiang, X. (2001). The Aspergillus cytoplasmic dynein heavy chain and NUDF localize to microtubule ends and affect microtubule dynamics. Curr Biol 11, 719–724.

19. Laan, L., Pavin, N., Husson, J., Romet-Lemonne, G., van Duijn, M., Lopez, M.P., Vale, R.D., Julicher, F., Reck-Peterson, S.L., and Dogterom, M. (2012). Cortical dynein controls microtubule dynamics to generate pulling forces that position microtubule asters. Cell 148, 502–514.

20. Coquelle, F.M., Caspi, M., Cordelieres, F.P., Dompierre, J.P., Dujardin, D.L., Koifman, C., Martin, P., Hoogenraad, C.C., Akhmanova, A., Galjart, N., et al. (2002). LIS1, CLIP-170’s key to the dynein/dynactin pathway. Mol Cell Biol 22, 3089–3102.

21. Grabham, P.W., Seale, G.E., Bennecib, M., Goldberg, D.J., and Vallee, R.B. (2007). Cytoplasmic dynein and LIS1 are required for microtubule advance during growth cone remodeling and fast axonal outgrowth. J Neurosci 27, 5823–5834.

22. Sapir, T., Elbaum, M., and Reiner, O. (1997). Reduction of microtubule catastrophe events by LIS1, platelet-activating factor acetylhydrolase subunit. EMBO J 16, 6977–6984.

23. Chiu, Y.H., Xiang, X., Dawe, A.L., and Morris, N.R. (1997). Deletion of nudC, a nuclear migration gene of Aspergillus nidulans, causes morphological and cell wall abnormalities and is lethal. Mol Biol Cell 8, 1735–1749.

24. Xiang, X., Osmani, A.H., Osmani, S.A., Xin, M., and Morris, N.R. (1995). NudF, a nuclear migration gene in Aspergillus nidulans, is similar to the human LIS-1 gene required for neuronal migration. Mol Biol Cell 6, 297–310.

25. Osmani, A.H., Osmani, S.A., and Morris, N.R. (1990). The molecular cloning and identification of a gene product specifically required for nuclear movement in Aspergillus nidulans. J Cell Biol 111, 543–551.

26. Aumais, J.P., Tunstead, J.R., McNeil, R.S., Schaar, B.T., McConnell, S.K., Lin, S.H., Clark, G.D., and Yu-Lee, L.Y. (2001). NudC associates with Lis1 and the dynein motor at the leading pole of neurons. J Neurosci 21, RC187.

27. Cappello, S., Monzo, P., and Vallee, R.B. (2011). NudC is required for interkinetic nuclear migration and neuronal migration during neocortical development. Dev Biol 357, 326–335.

28. Wynshaw-Boris, A., and Gambello, M.J. (2001). LIS1 and dynein motor function in neuronal migration and development. Genes Dev 15, 639–651.

29. Zheng, M., Cierpicki, T., Burdette, A.J., Utepbergenov, D., Janczyk, P.L., Derewenda, U., Stukenberg, P.T., Caldwell, K.A., and Derewenda, Z.S. (2011). Structural features and chaperone activity of the NudC protein family. J Mol Biol 409, 722–741.

30. Zhu, X.J., Liu, X., Jin, Q., Cai, Y., Yang, Y., and Zhou, T. (2010). The L279P mutation of nuclear distribution gene C (NudC) influences its chaperone activity and lissencephaly protein 1 (LIS1) stability. J Biol Chem 285, 29903–29910.

31. Drerup, C.M., and Nechiporuk, A.V. (2013). JNK-interacting protein 3 mediates the retrograde transport of activated c-Jun N-terminal kinase and lysosomes. PLoS Genet 9, e1003303.

32. Aumais, J.P., Williams, S.N., Luo, W., Nishino, M., Caldwell, K.A., Caldwell, G.A., Lin, S.H., and Yu-Lee, L.Y. (2003). Role for NudC, a dynein-associated nuclear movement protein, in mitosis and cytokinesis. J Cell Sci 116, 1991–2003.

33. Del Bene, F., Wehman, A.M., Link, B.A., and Baier, H. (2008). Regulation of neurogenesis by interkinetic nuclear migration through an apical-basal notch gradient. Cell 134, 1055–1065.

34. Drerup, C.M., Herbert, A.L., Monk, K.R., and Nechiporuk, A.V. (2017). Regulation of mitochondria-dynactin interaction and mitochondrial retrograde transport in axons. Elife 6.

35. Markus, S.M., Marzo, M.G., and McKenney, R.J. (2020). New insights into the mechanism of dynein motor regulation by lissencephaly-1. Elife 9.

36. Drerup, C.M., and Nechiporuk, A.V. (2016). In vivo analysis of axonal transport in zebrafish. Methods Cell Biol 131, 311–329.

37. Tan, S.C., Scherer, J., and Vallee, R.B. (2011). Recruitment of dynein to late endosomes and lysosomes through light intermediate chains. Mol Biol Cell 22, 467–477.

38. Song, Y., and Brady, S.T. (2015). Post-translational modifications of tubulin: pathways to functional diversity of microtubules. Trends Cell Biol 25, 125–136.

39. Jha, R., Roostalu, J., Cade, N.I., Trokter, M., and Surrey, T. (2017). Combinatorial regulation of the balance between dynein microtubule end accumulation and initiation of directed motility. EMBO J 36, 3387–3404.

40. Kardon, J.R., and Vale, R.D. (2009). Regulators of the cytoplasmic dynein motor. Nat Rev Mol Cell Biol 10, 854–865.

41. Duellberg, C., Trokter, M., Jha, R., Sen, I., Steinmetz, M.O., and Surrey, T. (2014). Reconstitution of a hierarchical +TIP interaction network controlling microtubule end tracking of dynein. Nat Cell Biol 16, 804–811.

42. Komarova, Y., De Groot, C.O., Grigoriev, I., Gouveia, S.M., Munteanu, E.L., Schober, J.M., Honnappa, S., Buey, R.M., Hoogenraad, C.C., Dogterom, M., et al. (2009). Mammalian end binding proteins control persistent microtubule growth. J Cell Biol 184, 691–706.

43. Hammond, J.W., Huang, C.F., Kaech, S., Jacobson, C., Banker, G., and Verhey, K.J. (2010). Posttranslational modifications of tubulin and the polarized transport of kinesin-1 in neurons. Mol Biol Cell 21, 572–583.

44. Hendershott, M.C., and Vale, R.D. (2014). Regulation of microtubule minus-end dynamics by CAMSAPs and Patronin. Proc Natl Acad Sci U S A 111, 5860–5865.

45. Baumbach, J., Murthy, A., McClintock, M.A., Dix, C.I., Zalyte, R., Hoang, H.T., and Bullock, S.L. (2017). Lissencephaly-1 is a context-dependent regulator of the human dynein complex. Elife 6.

46. Yamada, M., Toba, S., Takitoh, T., Yoshida, Y., Mori, D., Nakamura, T., Iwane, A.H., Yanagida, T., Imai, H., Yu-Lee, L.Y., et al. (2010). mNUDC is required for plus-end-directed transport of cytoplasmic dynein and dynactins by kinesin-1. EMBO J 29, 517–531.

47. Herbert, A.L., Fu, M.M., Drerup, C.M., Gray, R.S., Harty, B.L., Ackerman, S.D., O’Reilly-Pol, T., Johnson, S.L., Nechiporuk, A.V., Barres, B.A., et al. (2017). Dynein/dynactin is necessary for anterograde transport of Mbp mRNA in oligodendrocytes and for myelination in vivo. Proc Natl Acad Sci U S A 114, E9153–E9162.

48. Kevenaar, J.T., and Hoogenraad, C.C. (2015). The axonal cytoskeleton: from organization to function. Front Mol Neurosci 8, 44.

49. Dubey, J., Ratnakaran, N., and Koushika, S.P. (2015). Neurodegeneration and microtubule dynamics: death by a thousand cuts. Front Cell Neurosci 9, 343.

50. Hayashi, I., Wilde, A., Mal, T.K., and Ikura, M. (2005). Structural basis for the activation of microtubule assembly by the EB1 and p150Glued complex. Mol Cell 19, 449–460.

51. Honnappa, S., Okhrimenko, O., Jaussi, R., Jawhari, H., Jelesarov, I., Winkler, F.K., and Steinmetz, M.O. (2006). Key interaction modes of dynamic +TIP networks. Mol Cell 23, 663–671.

52. Lazarus, J.E., Moughamian, A.J., Tokito, M.K., and Holzbaur, E.L. (2013). Dynactin subunit p150(Glued) is a neuron-specific anti-catastrophe factor. PLoS Biol 11, e1001611.

53. Manna, T., Honnappa, S., Steinmetz, M.O., and Wilson, L. (2008). Suppression of microtubule dynamic instability by the +TIP protein EB1 and its modulation by the CAP-Gly domain of p150glued. Biochemistry 47, 779–786.

54. Adames, N.R., and Cooper, J.A. (2000). Microtubule interactions with the cell cortex causing nuclear movements in Saccharomyces cerevisiae. J Cell Biol 149, 863–874.

55. Kozlowski, C., Srayko, M., and Nedelec, F. (2007). Cortical microtubule contacts position the spindle in C. elegans embryos. Cell 129, 499–510.

56. Ligon, L.A., Karki, S., Tokito, M., and Holzbaur, E.L. (2001). Dynein binds to beta-catenin and may tether microtubules at adherens junctions. Nat Cell Biol 3, 913–917.

57. Heil-Chapdelaine, R.A., Oberle, J.R., and Cooper, J.A. (2000). The cortical protein Num1p is essential for dynein-dependent interactions of microtubules with the cortex. J Cell Biol 151, 1337–1344.

58. Ananthanarayanan, V., Schattat, M., Vogel, S.K., Krull, A., Pavin, N., and Tolic-Norrelykke, I.M. (2013). Dynein motion switches from diffusive to directed upon cortical anchoring. Cell 153, 1526–1536.

59. Gusnowski, E.M., and Srayko, M. (2011). Visualization of dynein-dependent microtubule gliding at the cell cortex: implications for spindle positioning. J Cell Biol 194, 377–386.

60. Perlson, E., Hendricks, A.G., Lazarus, J.E., Ben-Yaakov, K., Gradus, T., Tokito, M., and Holzbaur, E.L. (2013). Dynein interacts with the neural cell adhesion molecule (NCAM180) to tether dynamic microtubules and maintain synaptic density in cortical neurons. J Biol Chem 288, 27812–27824.

61. Nishino, M., Kurasawa, Y., Evans, R., Lin, S.H., Brinkley, B.R., and Yu-Lee, L.Y. (2006). NudC is required for Plk1 targeting to the kinetochore and chromosome congression. Curr Biol 16, 1414–1421.

62. Zhou, T., Aumais, J.P., Liu, X., Yu-Lee, L.Y., and Erikson, R.L. (2003). A role for Plk1 phosphorylation of NudC in cytokinesis. Dev Cell 5, 127–138.

63. Riera, J., and Lazo, P.S. (2009). The mammalian NudC-like genes: a family with functions other than regulating nuclear distribution. Cell Mol Life Sci 66, 2383–2390.

64. Glock, C., Biever, A., Tushev, G., Nassim-Assir, B., Kao, A., Bartnik, I., Tom Dieck, S., and Schuman, E.M. (2021). The translatome of neuronal cell bodies, dendrites, and axons. Proc Natl Acad Sci U S A 118.

65. Zhang, K., Foster, H.E., Rondelet, A., Lacey, S.E., Bahi-Buisson, N., Bird, A.W., and Carter, A.P. (2017). Cryo-EM Reveals How Human Cytoplasmic Dynein Is Auto-inhibited and Activated. Cell 169, 1303–1314 e1318.

66. Schlager, M.A., Hoang, H.T., Urnavicius, L., Bullock, S.L., and Carter, A.P. (2014). In vitro reconstitution of a highly processive recombinant human dynein complex. EMBO J 33, 1855–1868.

67. McKenney, R.J., Vershinin, M., Kunwar, A., Vallee, R.B., and Gross, S.P. (2010). LIS1 and NudE induce a persistent dynein force-producing state. Cell 141, 304–314.

68. Westerfield, M. (1993). The zebrafish book: a guide for the laboratory use of zebrafish (Brachydanio rerio), (Eugene, OR: M. Westerfield).

69. Kimmel, C.B., Ballard, W.W., Kimmel, S.R., Ullmann, B., and Schilling, T.F. (1995). Stages of embryonic development of the zebrafish. Dev Dyn 203, 253–310.

70. Miller, A.C., Obholzer, N.D., Shah, A.N., Megason, S.G., and Moens, C.B. (2013). RNA-seq-based mapping and candidate identification of mutations from forward genetic screens. Genome Res 23, 679–686.

71. Mo, W., and Nicolson, T. (2011). Both pre- and postsynaptic activity of Nsf prevents degeneration of hair-cell synapses. PLoS One 6, e27146.

72. Kwan, K.M., Fujimoto, E., Grabher, C., Mangum, B.D., Hardy, M.E., Campbell, D.S., Parant, J.M., Yost, H.J., Kanki, J.P., and Chien, C.B. (2007). The Tol2kit: a multisite gateway-based construction kit for Tol2 transposon transgenesis constructs. Dev Dyn 236, 3088–3099.

73. Lee, I.H., Cao, L., Mostoslavsky, R., Lombard, D.B., Liu, J., Bruns, N.E., Tsokos, M., Alt, F.W., and Finkel, T. (2008). A role for the NAD-dependent deacetylase Sirt1 in the regulation of autophagy. Proc Natl Acad Sci U S A 105, 3374–3379.

74. Sheets, L., He, X.J., Olt, J., Schreck, M., Petralia, R.S., Wang, Y.X., Zhang, Q., Beirl, A., Nicolson, T., Marcotti, W., et al. (2017). Enlargement of Ribbons in Zebrafish Hair Cells Increases Calcium Currents But Disrupts Afferent Spontaneous Activity and Timing of Stimulus Onset. J Neurosci 37, 6299–6313.

75. Zhang, Q., Li, S., Wong, H.C., He, X.J., Beirl, A., Petralia, R.S., Wang, Y.X., and Kindt, K.S. (2018). Synaptically silent sensory hair cells in zebrafish are recruited after damage. Nat Commun 9, 1388.

76. Obholzer, N., Wolfson, S., Trapani, J.G., Mo, W., Nechiporuk, A., Busch-Nentwich, E., Seiler, C., Sidi, S., Sollner, C., Duncan, R.N., et al. (2008). Vesicular glutamate transporter 3 is required for synaptic transmission in zebrafish hair cells. J Neurosci 28, 2110–2118.

77. Mandal, A., Wong, H.C., Pinter, K., Mosqueda, N., Beirl, A., Lomash, R.M., Won, S., Kindt, K.S., and Drerup, C.M. (2021). Retrograde Mitochondrial Transport Is Essential for Organelle Distribution and Health in Zebrafish Neurons. J Neurosci 41, 1371–1392.

78. Drerup, C.M., Lusk, S., and Nechiporuk, A. (2016). Kif1B Interacts with KBP to Promote Axon Elongation by Localizing a Microtubule Regulator to Growth Cones. J Neurosci 36, 7014–7026.

79. Schindelin, J., Arganda-Carreras, I., Frise, E., Kaynig, V., Longair, M., Pietzsch, T., Preibisch, S., Rueden, C., Saalfeld, S., Schmid, B., et al. (2012). Fiji: an open-source platform for biological-image analysis. Nat Methods 9, 676–682.

