## Supplemental Figures for "NudC regulated Lis1 stability is essential for maintenance of dynamic microtubule ends in the axon terminal"

Supplemental information

**S1.** The *nudc* mutation causes loss of NudC function and autophagosome accumulation in axon terminals. (**A**) Overexpression of mRFP-NudC in *nudc* mutants can suppress the axon terminal swelling phenotype (ANOVA). **(B)** RT-PCR for *nudc* shows it is maternally deposited (present at 2-cell stage) and also zygotically expressed. (**C,D**) Transmission electron micrographs of wild type and *nudc* axon terminals. Enlarged autophagosomes are labeled with yellow arrows. Axon terminal outlined by dashed line. **(E)** Quantification of autophagosome density in axons from TEM images (ANOVA). Wild type data was collected from 12 micrographs representing 3 fish. *nudc* data was collected from 9 micrographs representing 3 fish. Data are expressed as mean ± S.E.M.

**S2.** Mitochondrial and peroxisome transport in *nudc* mutants. **(A,B)** Kymograph analysis of peroxisomes (labeled by a peroxisome targeting sequence tagged with mRFP) transport. **(C)** Percent of peroxisomes in the anterograde, retrograde, stationary, and bidirectional populations are unaffected in *nudc* mutants. A slight, non-significant reduction in anterograde peroxisome transport was noted (ANOVA; p=0.0553). **(D)** Total number of peroxisomes present in the axon is unchanged in *nudc* mutants (ANOVA). **(E,F)** Quantification of peroxisome transport parameters reveals slight decreases in their anterograde distance and velocities in *nudc* mutants (ANOVA). **(F`,F``)** Histogram of velocities binned in 0.2μm/sec intervals demonstrates a slight shift towards slower anterograde velocities. **(G,H)** Kymograph analysis of mitochondrial transport. **(I)** Percent of mitochondria in the anterograde, retrograde, stationary and bidirectional populations are unaffected in *nudc* mutants (ANOVA). **(J)** Total number of mitochondria present in the axon is unchanged in *nudc* mutants (ANOVA). **(K,L)** Quantification of mitochondrial transport parameters reveals slight decreases in their anterograde and retrograde transport velocities in *nudc* mutants (Anterograde: Wilcoxon Rank Sum; Retrograde: ANOVA). **(L`,L``)** Histogram of velocities binned in 0.2μm/sec intervals demonstrates a shift towards slower velocities particularly in the retrograde direction. Data are expressed as mean ± S.E.M. Sample sizes indicated on graph.

**S3.** Dync1li1V2-labeled cargo transport in *nudc* mutants. **(A,B)** Kymograph analysis of Dync1li1v2 transport. **(C)** The percent of anterograde Dync1li1v2 transport decreases in *nudc* mutants (ANOVA; *p*=0.0002) while bidirectional movement increases (ANOVA; *p*=0.0009). Retrograde and stationary Dync1li1V2 punctal transport frequency are unchanged (ANOVA; *p*=0.3839 and *p*=0.069 respectively). **(D)** Total number of Dync1li1V2+ vesicles in the axon are unchanged in *nudc* mutants (ANOVA). **(E,F)** Quantification of Dync1li1v2 transport shows that anterograde and retrograde velocity are both reduced in *nudc* mutants (ANOVA). **(F`F``)** Shifts towards slower velocities are clear in the binned histograms. Data are expressed as mean ± S.E.M. Sample sizes indicated on graphs.

**S4.** Microtubule stability in the axon shaft is unchanged in *nudc* mutants. **(A,B)** Kymograph analysis of EB3-labeled microtubule growth. **(C)** The number of plus-end directed comets are the same between wild type and *nudc* mutants (ANOVA). **(D,E)** Quantification of microtubule growth distance and velocity show no difference between wild type and *nudc* mutants (ANOVA). **(D`, E`)** Binned histograms of distance and velocity showing no population shifts for either measurement. **(F,G)** Images of RFP-Patronin-CC labeling of microtubule minus ends in the cell body, axon, and axon terminal. **(H-J)** Quantification of Patronin punctal density shows no difference in microtubule minus end number in any compartment (ANOVA). Data are expressed as mean ± S.E.M. Scale bar = 10μm. Sample sizes indicated on graphs.

### Figure S1

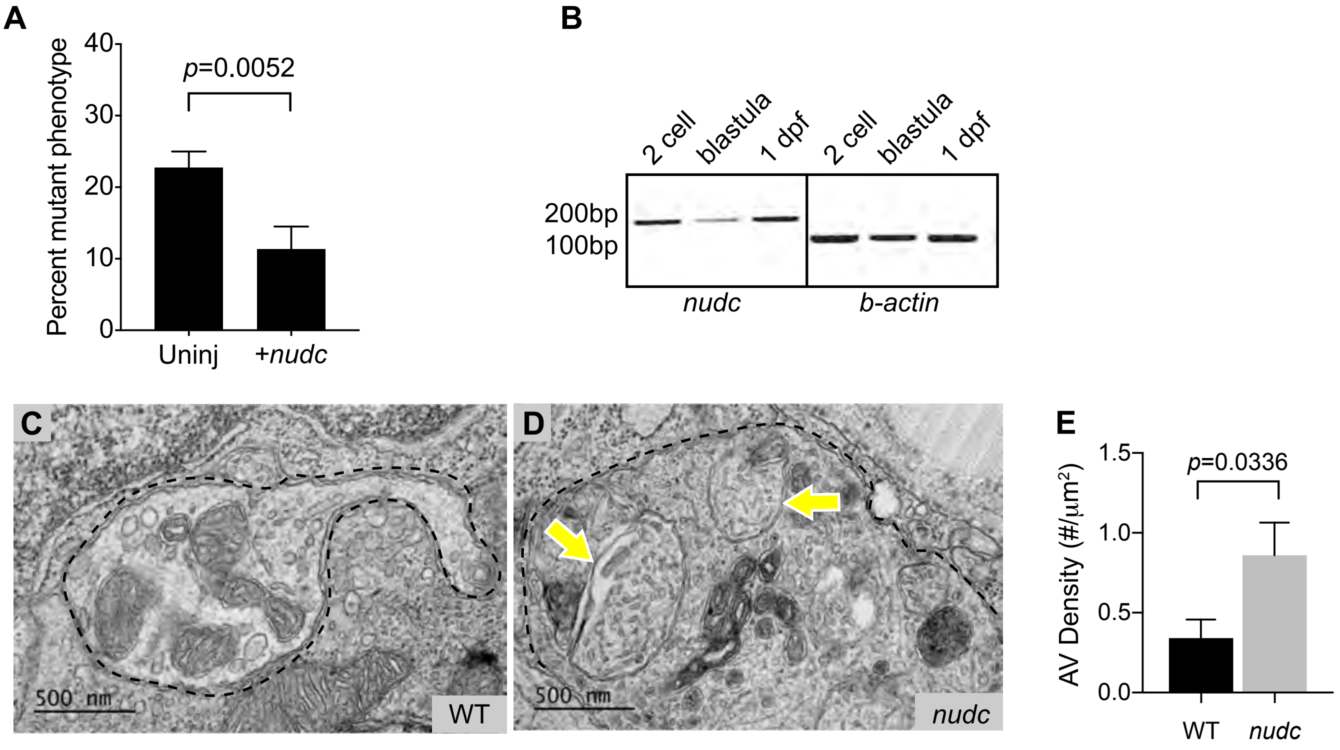

A
Percent mutant phenotype
40
30
20
10
0
p=0.0052
Uninj
+nudc
B
2 cell
blastula
1 dpf
2 cell
blastula
1 dpf
200bp
100bp
nudc
b-actin
C
500 nm
WT
D
500 nm
nudc
E
AV Density (#/μm²)
1.5
1.0
0.5
0.0
p=0.0336
WT
nudc

### Figure S2

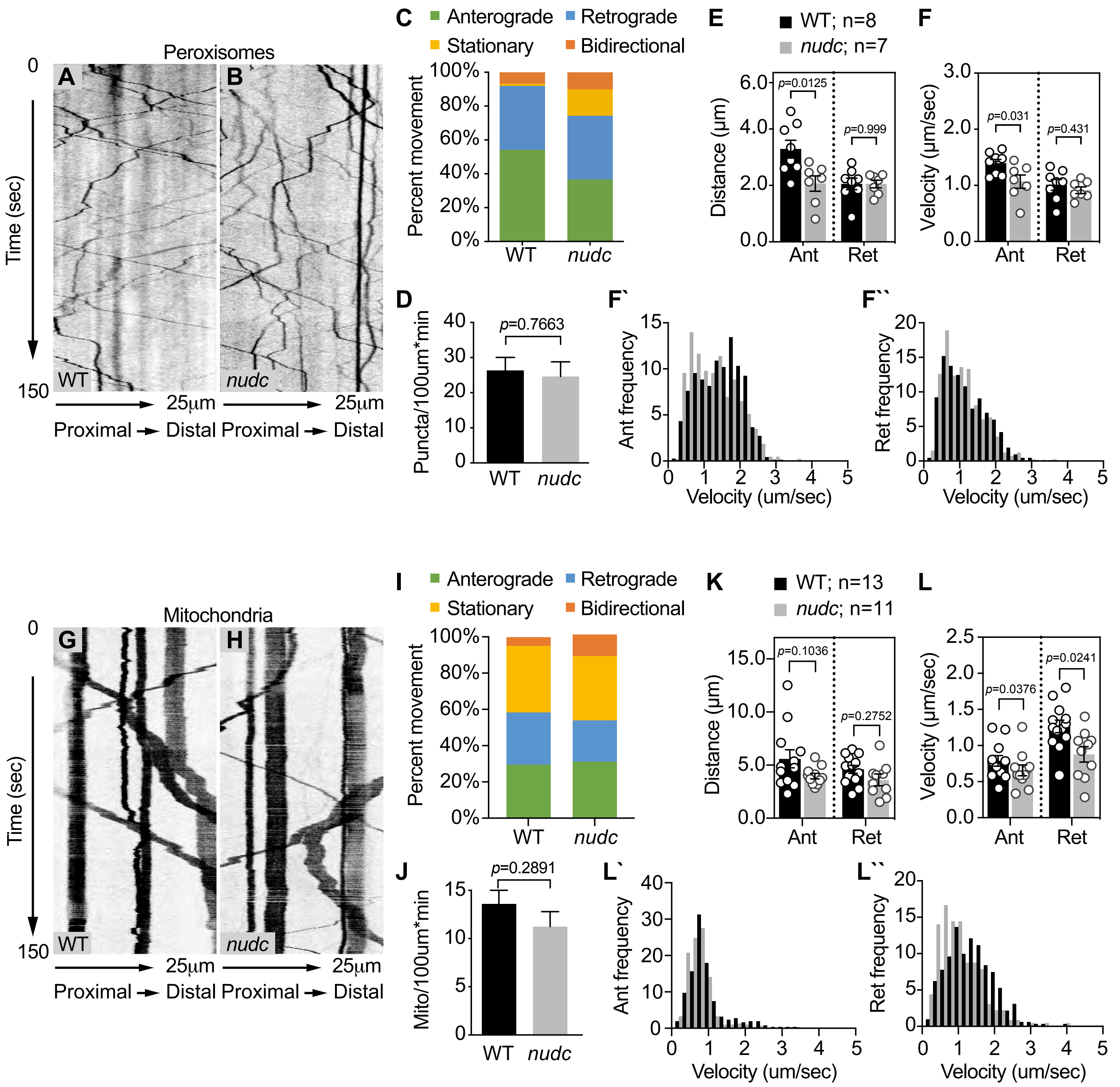

Peroxisomes
A
B
0
150
Time (sec)
WT
nudc
25µm
25µm
Proximal → Distal Proximal → Distal
C
Anterograde
Retrograde
Stationary
Bidirectional
Percent movement
100%
80%
60%
40%
20%
0%
WT
nudc
E
WT; n=8
nudc; n=7
Distance (µm)
6.0
4.0
2.0
0.0
p=0.0125
p=0.999
Ant
Ret
F
Velocity (um/sec)
3.0
2.0
1.0
0.0
p=0.031
p=0.431
Ant
Ret
D
Puncta/100um*min
40
30
20
10
0
p=0.7663
WT
nudc
F`
Ant frequency
15
10
5
0
0 1 2 3 4 5
Velocity (um/sec)
F``
Ret frequency
20
15
10
5
0
0 1 2 3 4 5
Velocity (um/sec)
Mitochondria
G
H
0
150
Time (sec)
WT
nudc
25µm
25µm
Proximal → Distal Proximal → Distal
I
Anterograde
Retrograde
Stationary
Bidirectional
Percent movement
100%
80%
60%
40%
20%
0%
WT
nudc
K
WT; n=13
nudc; n=11
Distance (µm)
15.0
10.0
5.0
0.0
p=0.1036
p=0.2752
Ant
Ret
L
Velocity (um/sec)
2.5
2.0
1.5
1.0
0.5
0.0
p=0.0376
p=0.0241
Ant
Ret
J
Mito/100um*min
15
10
5
0
p=0.2891
WT
nudc
L`
Ant frequency
40
30
20
10
0
0 1 2 3 4 5
Velocity (um/sec)
L``
Ret frequency
20
15
10
5
0
0 1 2 3 4 5
Velocity (um/sec)

### Figure S3

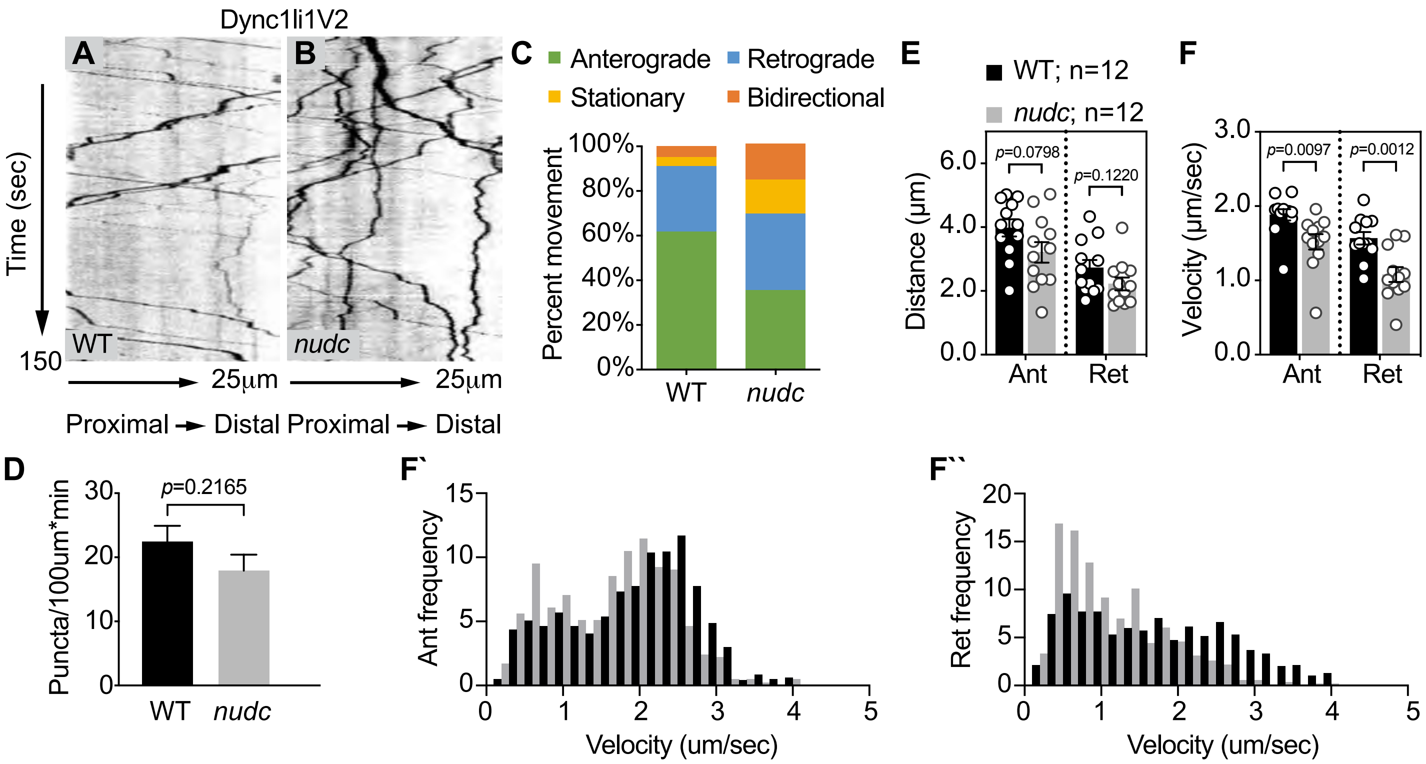

Dync1li1V2
A
B
Time (sec)
150
WT
nudc
25µm
25µm
Proximal → Distal Proximal → Distal
C
Anterograde
Retrograde
Stationary
Bidirectional
Percent movement
100%
80%
60%
40%
20%
0%
WT
nudc
E
WT; n=12
nudc; n=12
Distance (µm)
6.0
4.0
2.0
0.0
p=0.0798
p=0.1220
Ant
Ret
F
Velocity (µm/sec)
3.0
2.0
1.0
0.0
p=0.0097
p=0.0012
Ant
Ret
D
Puncta/100um*min
30
20
10
0
p=0.2165
WT
nudc
F`
Ant frequency
15
10
5
0
0 1 2 3 4 5
Velocity (um/sec)
F``
Ret frequency
20
15
10
5
0
0 1 2 3 4 5
Velocity (um/sec)

### Figure S4

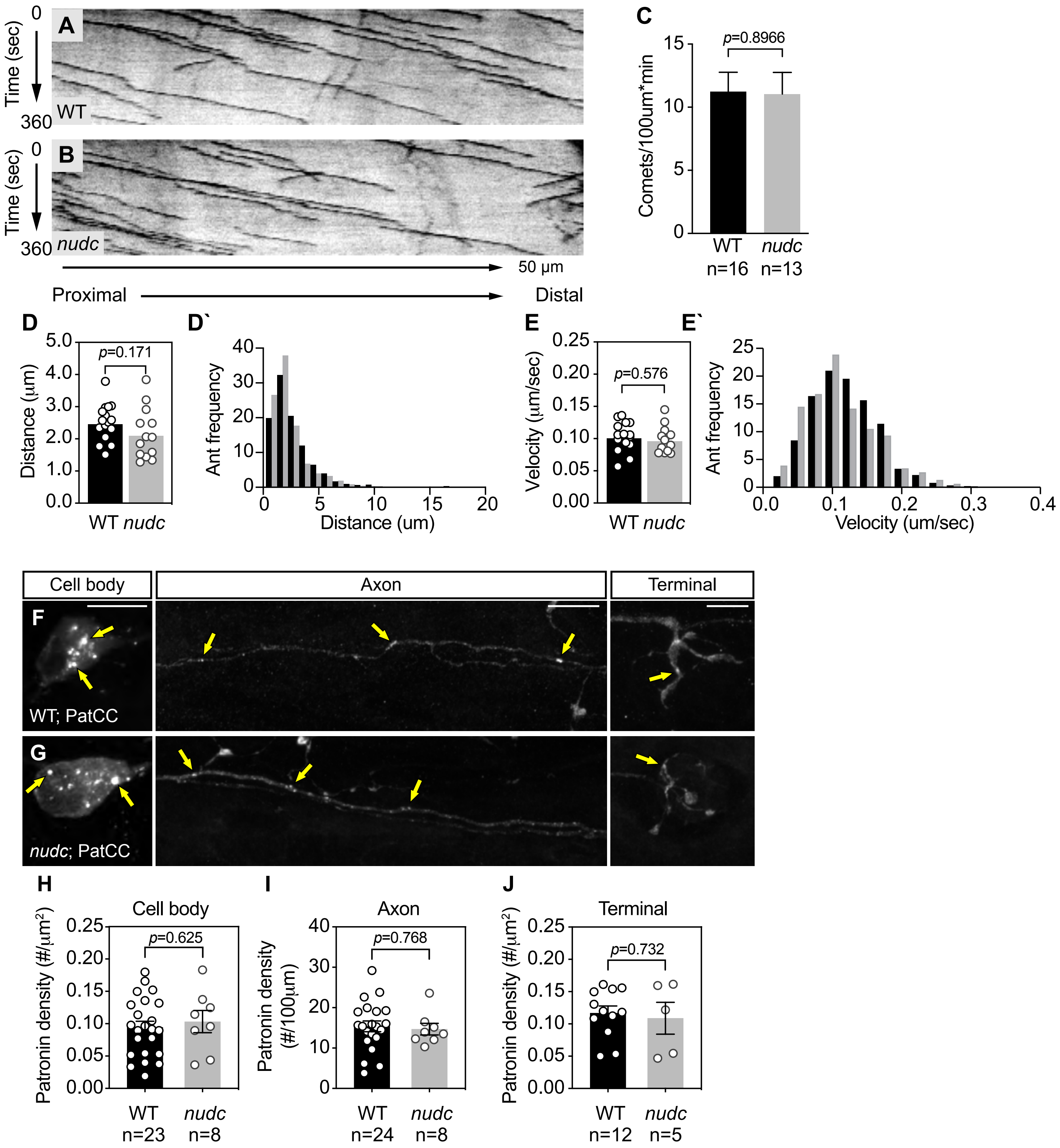

A
B
C
D
D`
E
E`
F
G
H
I
J
Time (sec)
0
360
WT
nudc
50 μm
Proximal
Distal
Comets/100um*min
p=0.8966
WT n=16
nudc n=13
Distance (μm)
p=0.171
Ant frequency
Distance (um)
Velocity (μm/sec)
p=0.576
Velocity (um/sec)
Cell body
Axon
Terminal
WT; PatCC
nudc; PatCC
Patronin density (#/μm²)
Patronin density (#/100μm)
p=0.625
p=0.768
p=0.732
WT n=23
nudc n=8
WT n=24
nudc n=8
WT n=12
nudc n=5
